# Autophagy slows the aging of Germline stem cells in *Drosophila* through modulation of E-cadherin

**DOI:** 10.1101/2022.03.31.486570

**Authors:** Nidhi Murmu, Bhupendra V. Shravage

**Affiliations:** Developmental Biology Group, MACS-Agharkar Research Institute, G G Agarkar Road, Pune −411004; Department of Biotechnology, Savitribai Phule Pune University, Ganeshkhind, Pune-411004

**Keywords:** Autophagy, germline stem cells, aging, autophagy flux, E-cadherin

## Abstract

Autophagy is a conserved process that degrades cytoplasmic components and organelles in metazoan cells including germline stem cells. Although autophagy is implicated in the aging of stem cells, the precise mechanism are still unknown. Here we show that elevating autophagy by overexpressing (OE) *Drosophila* Autophagy-related gene 8a (Atg8a) in the female Germline stem cells (GSCs) delays their loss due to aging. However, sustained elevated autophagy levels in old flies promote GSC loss due to cell death. In contrast, knockdown of Atg8a (*Atg8aRNAi*) in GSCs accelerates their loss. Atg8a^OE^ GSCs show elevated autophagy flux, and increased mitotic activity even at 8 weeks of age. Atg8a^OE^ GSCs possess smaller-sized mitochondria and exhibit reduced mitochondrial oxidative stress in the GSCs. However, in contrast *Atg8aRNAi* GSCs have elevated mitochondrial ROS and possess larger mitochondria. Finally, our data show that Atg8a^OE^ GSCs occupy the stem cell niche for longer duration with the aid of elevated E-cadherin at the GSC-cap cell contact sites. Our data suggests that elevated autophagy promotes GSC maintenance and activity, and delays their aging.

**Graphical abstract:** 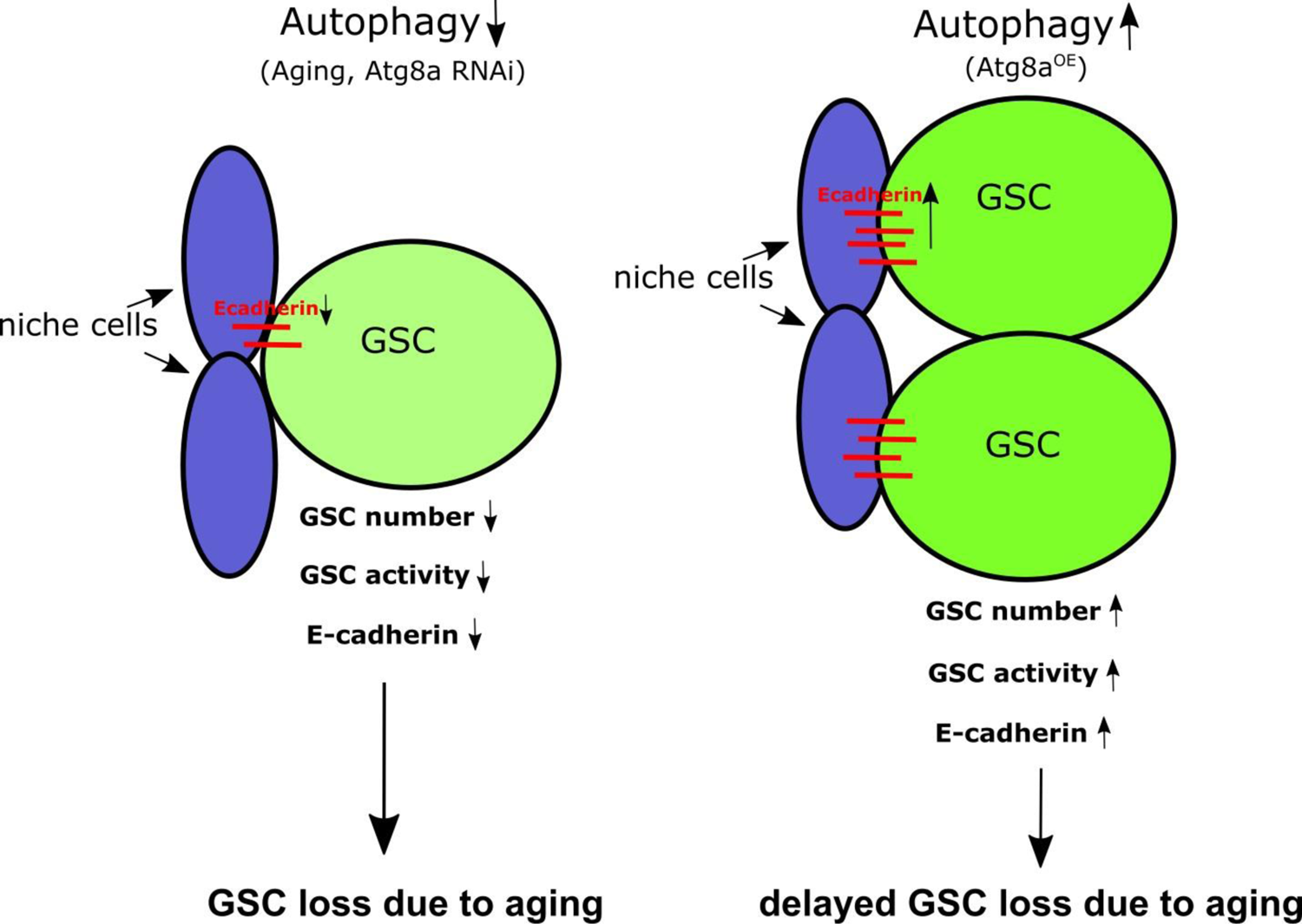

## Introduction

Germline stem cells (GSCs) are specialized cells that divide asymmetrically to give rise to a daughter GSC and cystoblast (Fuller and Spradling, 2007). This activity is important for the constant replenishment of stem cell pools affecting tissue homeostasis. Specific microenvironments called ‘niche’ maintain these stem cells by regulating signals that modulate their activity(Morrison and Spradling, 2008). Although stem cells are thought to divide infinitely throughout their lifetime, these cells also fall prey to the consequences of aging due to the accumulation of cellular damage (Liu and Rando, 2011; Oh et al., 2014; Phadwal et al., 2013; Signer and Morrison, 2013). Germline stem cell number and division rate progressively decline as a function of age. This decline is attributed to reduced expression of regulatory signals from the niche. The constant self-renewal of germline stem cells demands activation of mechanisms that mediate the removal of cellular damage (Ishibashi et al., 2020; Tolkin and Hubbard, 2021). Autophagy is one such mechanism that plays a key role in clearing out cellular damage by degrading protein aggregates and damaged organelles via delivery to lysosomes. However, it remains unclear if autophagy influences germline stem cell maintenance and aging.

Macroautophagy (autophagy) is an essential process operational within all metazoan cells that involves degradation of toxic proteins and damaged organelles through the lysosomal system. It is characterized by the formation of double membrane vesicle called autophagosome that carries the cargo for degradation. Fusion of autophagosome with the lysosome forms the autolysosome wherein degradation of the cargo occurs and the degradation products are recycled back in to the cytoplasm for use in cellular processes (Mizushima, 2007). Autophagy is a complex process governed by several Autophagy-related (Atg) proteins. 18 Atg proteins form 5 complexes that drive the process of autophagy in *Drosophila* (Lőrincz et al., 2017; Mulakkal et al., 2014). The ubiquitin like protein Atg8 (Atg8a and Atg8ab in *Drosophila*, LC3 family protein in higher organism) is crucial for autophagosome formation, elongation and closure. Atg8a mutants in *Drosophila* are devoid of autophagosomes in several tissues (Simonsen et al., 2008). Atg8a is mutants have defects in several tissues and die early as compared to the control flies. Reduced Atg8a leads to a decrease in autophagy flux. Autophagy flux is the amount and the rate at which cargo is degraded in the lysosomes. Upon upregulation of autophagy there is an increase in degradation of cargo. Autophagy flux declines with aging in cells and is partly responsible for loss of proteostasis (Aman et al., 2021; López-Otín et al., 2013; Oh et al., 2014).

Autophagy is essential to maintain a healthy life span in *Drosophila*. Several Atg mutants in *Drosophila* exhibit signs of accelerated aging. For instance, Atg7 and Atg8a mutants in *Drosophila* have impaired autophagy leading to accumulation of protein aggregates in neurons leading to early death (Juhasz et al., 2007; Simonsen et al., 2008). Further these mutants have reduced tolerance to stress such as nutrient stress and oxidative stress. Studies from several different model organisms including mouse models have revealed that elevating autophagy supports cellular homeostasis leading to extension of lifespan. For instance, neuronal expression of Atg8a and Atg1 in *Drosophila* leads to upto 50% increase in lifespan over control animals. Moderate Atg1 expression in fatbody, intestine and Malpighian tubules in *Drosophila* leads to extension of lifespan through modulation of mitochondrial genes and improved proteostasis (Bjedov et al., 2020). Additionally, elevated levels of autophagy cargo receptor Ref(2)P (p62 in mammals) also improved proteostasis, facilitated mitophagy and enhanced mitochondrial function (Aparicio et al., 2019). Enhancing mitophagy improves mitochondrial health and homeostasis which subsequently translates to extension of life span through activation of pro-longevity metabolism (Rana et al., 2017).

An important function of autophagy is to alleviate generation and mitigation of reactive oxygen species (ROS) generated during metabolic reactions (Scherz-Shouval and Elazar, 2011). Mitochondria are the primary sources of ROS such as 0_2_, H_2_O_2_ and OH (Shadel and Horvath, 2015). These ROS have potential to damage macromolecular proteins, DNA and lipids via oxidation which subsequently negatively affects ATP production. Mitochondrial ROS (mROS) in low amounts stimulates autophagy and in higher amounts triggers mitophagy(Zhou et al., 2021). Removal of damaged mitochondria through the process of mitophagy is critical for reducing ROS levels within cells and achieving redox homeostasis. ROS are also neutralized by several scavengers present within the cytoplasm and mitochondria(Dan Dunn et al., 2015; Shadel and Horvath, 2015). Redoxins such as glutaredoxins and peroxiredoxins function in non-enzymatic pathways to maintain reduced state by themselves getting oxidized and in process transfer the electrons to glutathione. Enzymes such as catalase, superoxide dismutase neutralize ROS enzymatically (Chen et al., 2009; Dröge, 2002). Inability or reduced ability to mitigate ROS is also responsible for accelerating aging of cells (López-Otín et al., 2013).

The female germline stem cells in *Drosophila* ovary are an ideal system to investigate the precise role of autophagy in stem cell maintenance and aging (He and Jasper, 2014). Each fly ovary is made up of 16-20 ovarioles that function to produce eggs. The proximal tip of the ovariole is called germarium that harbors stem cells necessary for generating the egg (egg chamber in *Drosophila*). Two to three germline stem cells reside in a well characterized niche made up by terminal filament cells, cap cells and escort stem cells. The signaling pathways controlling GSC self-renewal are well studied (Fuller and Spradling, 2007; Spradling., 1993). For instance, Dpp (BMP2/4 orthologue) is secreted from the cap cells and maintains GSCs. Loss of Dpp leads to loss of GSCs from the niche while constitutively active Dpp signaling leads to tumor like stem cell phenotype (Xie and Spradling, 1998). Most importantly, loss of GSC from the niche due to aging has been shown to be partly due to decrease in Dpp self-renewal signal (Pan et al., 2007; Zhao et al., 2008). Another molecule which is crucial for GSC maintenance is E-cadherin. E-cadherin is a component of adherens junction that adheres the GSC to the cap cells (Song et al., 2002). E-cadherin expression has been shown to reduce with age and this causes loss of GSCs from the niche and upregulating E-cadherin in GSCs leads to their longer retention in the niche (Pan et al., 2007; Zhao et al., 2008). Together, Dpp and E-cadherin maintain GSCs by regulating self-renewal and niche occupancy respectively.

In this study, we have addressed the role of autophagy in aging of female germline stem cells in *Drosophila*. There is an age-related decline in autophagy flux in GSCs. Blocking autophagy accelerated GSC loss with increased mitochondrial ROS generation and reduced E-cadherin expression. We investigated if elevating autophagy flux through Atg8a overexpression (Atg8a^OE^) would slow aging of GSCs. Our data show that Atg8a^OE^ leads to an increase in autophagy flux. It leads to longer retention of actively dividing GSCs in the niche. Further, our data show that Atg8a^OE^ promotes redox homeostasis. Finally we show that Atg8a^OE^ modulates E-cadherin expression at the GSC-niche boundary site and thereby facilitates niche occupancy of GSCs for longer duration.

## Materials and Methods

### Fly maintenance

Flies were maintained at 25 (degree) celsius with 12 hour day/light cycle.

### Fly strains

*OregonR*, *ywhsFLP1; mCherry-Atg8a/CyO; GFP-Ref(2)P/TM6b (Nilangekar et al., 2019) nanos-mito-roGFP2-Grx1 (Nilangekar and Shravage, 2019). The following flies were obtained from Bloomington Drosophila Stock Center, USA; yw; UASp.mCherry.Atg8a; Dr/ TM3, Ser (BL37750, RRID:BDSC_37750), w; +; nosGal4VP16 (BL4937, RRID:BDSC_4937).* The following RNAi lines: *y sc v; +; Atg8a-RNAi (BL34340, RRID:BDSC_34340)* and *y sc v; Atg8a-RNAi;+ (BL58309, RRID:BDSC_58309)* were combined by crossing to obtain *ywhsFLP1/12; Atg8a RNAi; Atg8a RNAi*.

### Fly food

Flies were reared on food of the following composition; 8 % sugar, 7.5 % corn flour, 3 % malt extract, 3 % yeast and 1 % agar. The food also contained anti-fungal chemicals; 0.12 % methyl benzoate, 0.08 % orthophosphoric acid and 0.4 % propionic acid.

### Chloroquine treatment

Stock solution of 50 mg/ml chloroquine in water was added during food preparation to a final concentration of 3 mg/ml when the food cooled down to 60°C. The flies were transferred to chloroquine containing food every day for two days.

### Aging of flies

For the experimental set-up, three different timepoints were studied; 1 week (young), 4 weeks (mid-aged) and 8 weeks (old). 250 to 300 flies of the desired genotype were collected for each time-point in fly food vials such that each vial housed not more than 15 females and 10 males. The collection of the flies were synchronized such that they attain the age of 1 week, 4 weeks and 8 weeks on the same day. Aging was also set-up wherein 250-300 flies were allowed to age and at the designated intervals 15-20 females were dissected for assays described. Flies were transferred to fresh food vials containing dry yeast pellets every 3^rd^ day till they reached the required age. We did not find significant differences in our assays in the different experimental set up of aging.

### Ovary dissection

Flies were dissected as described in (Nilangekar et al., 2019). For RNA extraction, ovaries were dissected in Grace’s medium and transferred to a 1.5 ml microfuge tube containing TriZol (Invitrogen, USA) and frozen at −80 C till further processing.

### Immunostaining

Ovaries were fixed in 300 μl of 4 % paraformaldehyde in 1xPBS, pH 7.4 for 20 minutes at room temperature. Washed thrice with 300 μl of 0.1 % PBTx (0.1 % Triton-x-100 in 1xPBS, pH 7.4). Blocked in 1% PBTx containing 0.5 % BSA (bovine serum albumin) for one hour at room temperature. Primary antibody was diluted in 0.3 % PBTx containing 0.5 % BSA to a total solution volume of 100-150 μl and incubated at 4 ᵒC overnight with 5 RPM rotation speed. The following day, the primary antibody was washed off, with 300 μl 0.1 % PBTx for 15 minutes at room temperature. The tissue was again blocked in 300 μl 0.1 % PBTx containing 10 % NGS (normal goat serum) for two hours at room temperature followed by secondary antibody diluted in 250 μl 0.1 % PBTx containing 10 % NGS for two hours at room temperature, from this onwards, the ovaries were protected from exposure to light. The secondary antibody was washed off with three washes of 300 μl 0.1 % PBTx for 15 minutes each at room temperature. The ovaries were then stained with 1 μg/ml DAPI in 300 μl 0.1 % PBTx for 10 min followed by two washes of 0.1 % PBTx for five minutes each at room temperature. The solution was removed and replaced with mounting medium; 1:1 ProLong Gold (Invitrogen, USA).

The following primary antibodies and dilutions were used; anti-CathepsinL 1:400 (Abcam Cat# ab58991, RRID:AB_940826), anti-ATP5A1 1:250 (Thermo Fisher Scientific Cat# 43-9800, RRID:AB_2533548), anti-E-cadherin 1:20 (DSHB Cat# DCAD2, RRID: AB_528120), anti-SMAD3 1:100 (Abcam Cat# ab52903, RRID: AB_882596), anti-Alpha spectrin 1:10 {DSHB Cat# 3A9(323/H10-2), RRID: AB_528473}, anti-hts 1B1 1:50 (DSHB Cat# 1B1-s), anti-LaminC 1:50 (DSHB Cat# LC28.26, RRID: AB_528339), anti-Caspase 1:200 (Abcam Cat# ab13847, RRID: 443014).

Secondary antibodies used: Alexa fluor 488 goat anti-rabbit 1:250 (Thermo Fisher Scientific Cat# A-11034, RRID:AB_2576217), Alexa fluor 555 goat anti-rat 1:250 (Thermo Fisher Scientific Cat# A-21434, RRID:AB_ 2535855), Alexa fluor 647 goat anti-mouse 1:250 (Thermo Fisher Scientific Cat# A-21236, RRID:AB_2535805), Alexa fluor 647 goat anti-rabbit 1:250 (Thermo Fisher Scientific Cat# A-21245, RRID:AB_2535813).

### DHE staining protocol

5-8 pair of ovaries were dissected in Grace’s medium (GM) followed by washing with 300ul GM for 4min. 30µM DHE in 300ul GM was added (protected from light) and incubated for 20 min at room temperature. The dye was washed off with one wash of GM for 5 min. The tissue was then fixed in 4% PFA for 10 min. The sample was incubated in 1 μg/ml DAPI solution in 300 μl 0.1 % PBTx for 5 minutes. Final washes of 0.1 % PBTx for five minutes each were given at room temperature.

### RNA isolation

35 pair of ovaries were used for RNA extraction using TriZol method (Invitrogen, USA). RNA extraction was carried out as per manufacturer’s instructions. The aqueous phase obtained after phase separation was then transferred to a fresh microfuge tube followed by addition of equal volumes of 95% ethanol and mixed thoroughly to precipitate the RNA. Further steps were performed using Monarch Total RNA Miniprep Kit (NEB #T2010). RNA was eluted in 20μl DEPC treated water.

### cDNA synthesis

1 µg of tissue specific (ovary) RNA was used for First strand cDNA synthesis using PrimeScript^TM^ 1^st^ strand cDNA Synthesis Kit (Takara, Cat# 6110A) using the following program: 65 °C for 5 min followed by 40 °C for 60 min. The synthesized cDNA was diluted (1:7.5) and validated with the help of PCR using reference genes at different cycles (25,30,32). The cDNA samples were then stored at −20 °C.

### Real-time PCR

qPCR was performed on EcoMax (Bibby Scientific, UK). Reactions were run in triplicates in a 48-well plate with 10 µl reaction volume. Following is the program used: 95°C for 10 minutes, 95°C for 15 seconds, 60° for 1 minute, 70°C for 5 seconds, 70°C for 5 seconds, 90°C for 2 minutes. The mean of housekeeping gene Tubulin was used as an internal control for normalization. The expression data was analysed using 2^-ΔΔCT^ described by (Livak and Schmittgen, 2001).

### Imaging and analysis of Mito-roGFP2-Grx1

The image acquisition and analysis was performed as mentioned in (Nilangekar and Shravage, 2019; Nilangekar et al., 2019)

### Imaging and Analysis

Leica SP8 Confocal microscope was used for imaging using 63x oil objective. 8-bit images were acquired at 100Hz scanning with pixel resolution of 1024 × 1024. Frame accumulation was performed as per the requirement of the channel. Analysis of confocal images was performed using ImageJ software. For puncta count, firstly, background subtraction was carried out using rolling ball plugin. The thresholding was performed objectively in a way that majority of the intense signal is distinguished from the background. A manual ROI was drawn around the germarium and GSC followed by particle analysis (size – size range, circularity 0.00-1.00 showing outlines). A resultant window displaying the particle size distribution of the corresponding threshold was obtained which was further evaluated in Microsoft Excel.

### Statistical analysis

Statistical analysis was performed using Student’s *T*-test for two samples assuming unequal variance for multiple comparisons using Microsoft Excel. The differences between two groups were considered significant at P<0.05. All the graphs were plotted in GraphPad Prism 7.

### Measurement of autophagic flux

To monitor flux, we have used reporters-nosP mCherry-Atg8a nos 3’UTR and nosP-GFP-Ref(2)P nos 3’UTR previously characterized in our lab ((Nilangekar et al., 2019)). The vesicles were identified by following the fluorescent markers and antibodies which has been sequentially explained. Both autophagosome and autolysosome membrane (briefly) consists of Atg8a protein. CathepsinL is a cysteine protease which marks the lysosome. Lysosome and autolysosomes consist of the same acid proteases, both the vesicles can be observed by CathepsinL. Since mCherry-Atg8a (autophagosomes) fuse with lysosome (CathepsinL), all resultant autolysosomal vesicles are mCherry-Atg8a-CathepsinL colocalized puncta. As the acidic pH of the autolysosomes quenches the GFP of GFP tagged Ref(2)P, GFP-Ref(2)P-CathepsinL colocalization signifies only the fusion event between the cargo and autolysosomes. The identification of these vesicles has been summarized in the following table.

| Marker used | Identification of the vesicles |
| --- | --- |
| mCherry-Atg8a alone | Autophagosome + Autolysosome |
| GFP-Ref(2)P alone | Cargo |
| CathepsinL alone | Autolysosome + lysosome |
| mCherry-Atg8a-CathepsinL | Autolysosome |
| GFP-Ref(2)P- CathepsinL | Fusion events between cargo and autolysosome |

### Measurement of mitochondrial area and number

The mitochondrial number and area were measured using Mitochondria Analyzer plugin in ImageJ software. The following operations were performed for mitochondrial analysis: Open ImageJ → Plugins→ Mitochondria Analyzer→ 2D→ 2D Threshold. In the resultant window, following values were fed for the specified parameters: Max Slope-1.5, Gamma −1.2, Block size-2.6, other values were kept default. The resultant thresholded image was saved and used for analysis.

### Analysis of mitochondrial area and number

A region of interest (ROI) was drawn using Freehand selections in the thresholded image, ROI was saved in ROI manager (Analyze →Tools → ROI Manager). The image with the drawn ROI was selected and following operations were performed: Plugins→ Mitochondria Analyzer→ 2D→ 2D Analysis. In the output window, analysis was performed on a per-mito basis. The resultant window was saved and the values were used to plot graphs.

## Results

### Autophagy flux reduces significantly in the germanium with age

Loss of proteostasis is one of the major hallmarks of aging in metazoans (López-Otín et al., 2013). Protein degradation machinery including autophagy within the cells is crucial for maintaining proteostasis. Several studies have shown that decline in autophagy contributes to aging (Aman et al., 2021). So we sought to test if autophagy is reduced in germline cells including GSCs during oogenesis as the flies age. To monitor autophagy we followed the expression of two candidates Atg8a fused to mCherry (LC3/GABARAP in mammals) and Ref(2)P fused to GFP (p62 in mammals) for our study. Atg8a is a ubiquitin-like protein that labels the inner and outer membranes of all autophagic structures: phagophore, autophagosomes, and autolysosomes, and has been widely used to monitor autophagy (Bali and Shravage, 2017; Klionsky et al., 2021; Nilangekar et al., 2019; Scott et al., 2004). Similarly, Ref(2) P (p62 in mammals), is a selective autophagy cargo receptor that binds polyubiquitinated substrates via ubiquitin-binding domain and interacts with Atg8a with the help of an Atg8-interacting motif (AIM/LIR) to recruit substrates to autophagosomes (Klionsky et al., 2021; Nezis et al., 2008). In order to monitor autophagy in the germline, we used germline-specific fluorescent reporter lines nosP-mCherry-Atg8a-nos3’UTR and nosP-GFP-Ref(2)P-nos3’UTR (GFP-Ref(2)P previously generated in our lab. We constructed a dual transgenic reporter by combining these transgenes. This double transgenic line was aged and the ovarian tissue harvested at 1 week, 4week and 8weeks of age and stained with lysosomal marker CathepsinL (a cysteine proteinase) to enable measurement of autophagy flux (Nilangekar et al., 2019).

Previous studies and data from our group have shown that basal autophagy occurs at very low levels in the GSCs, while significantly higher levels of autophagy could be detected in region 2 of the germarium (Nilangekar et al., 2019; Zhao et al., 2018). We decided to measure autophagy flux in both GSCs and germarium during aging. Autophagy flux is a measure of the degradation of cargo within the autolysosome. In the assays performed in this study, we measure autophagy flux by analyzing the autophagosomes and autolysosomes in both germarium and GSCs (for details refer to materials and methods). The higher the number of autophagosomes (AP) and autolysosomes (AL), the higher the autophagy flux. At 1 week old germaria and GSCs, the basal autophagy flux was relatively high. At 4 weeks, our analysis showed a significant decline in ALs in the germarium and both APs and ALs in GSCs. As compared to 1 week, we observed an overall decrease in both these vesicles in 8-week old germaria and GSCs (**Figure 1A-D’**). An interesting observation was an increase in AP and AL number from 4 to 8 weeks specifically in the GSCs (**Figure 1D-D’ and Supplementary Figure 1A-C”).** We also observed an accumulation of cargo receptor GFP-Ref(2)P due to reduced AP and AL formation (**Supplementary Figure 1B-B”**). Taken together, our data suggests that with age there is significant decrease in the formation of both autophagosomes and autolysosomes resulting in decreased autophagy flux in GSCs and germline cells.

**Figure 1.**
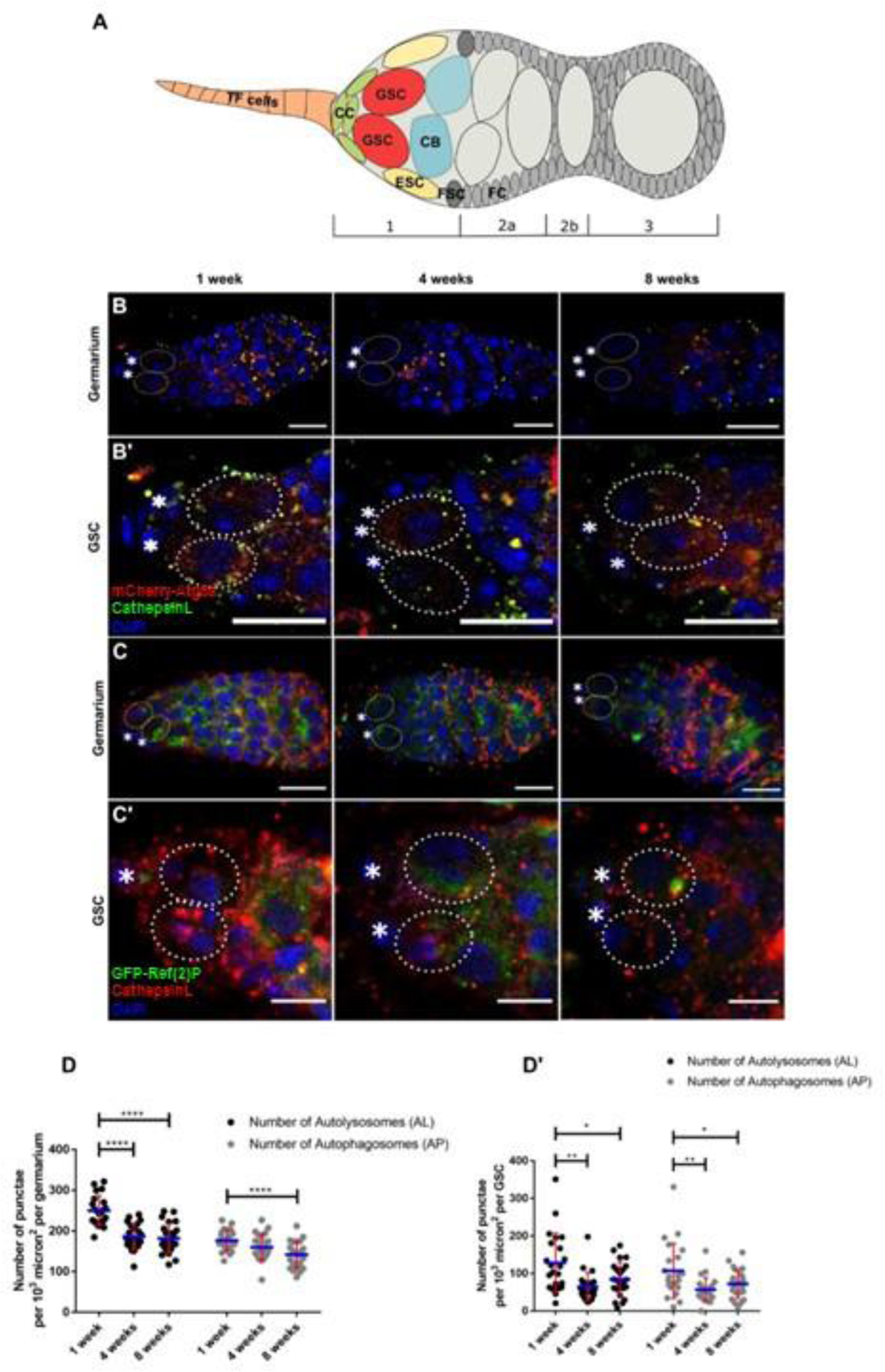
Autophagic flux reduces with age. (A) Schematic of a germarium. Germline stem cells (GSCs) reside at the anterior tip of the germarium. Cap cells (CC), terminal filament cells (TF) and escort stem cells (ESC) form niche to the GSCs. Abbreviations: FSC-Follicle stem cells, CB-Cystoblast. FC-Follicle cells. (B-C’) Flies of wildtype germaria (ywhsFLP1; nosP mCherry Atg8a; nosP GFP Ref(2)P) were aged at 1 week, 4 weeks and 8 weeks and stained for anti-CathepsinL to measure colocalization between mCherry Atg8a-CathepsinL in germarium (B) (scale bar-10µm) and GSCs (B’) (scale bar-5µm), GFP Ref(2)P-CathepsinL in germarium (C) (scale bar-10µm) and GSCs (C’) (scale bar-5µm), all marked in yellow (autolysosomes). Nuclei are marked in blue. Dotted ovals mark the GSCs and asterisk mark the cap cells. (D-D’) Interleaved scatter graph representing overall decrease in autolysosomes and autophagosomes in germarium (D) and GSCs (D’) respectively across 1 to 8 weeks. Error bars represent standard deviation (S.D.) in red and blue represent mean. n=20, Genotype: ywhsFLP1; nos mCherryAtg8a/+; nos GFP Ref(2)P/+. *p < 0.05, **p < 0.01, ****p < 0.0001

### Reducing autophagy increases ROS, accelerates GSC loss and leads to decrease in E-cadherin levels at the GSC-niche cell contact sites

Our data suggests that aging leads to decrease in autophagy flux in germline cells during oogenesis. Previous work from several groups have shown that aging leads to decline in GSC number and activity. This decline is due to reduced GSC-self renewal, niche occupancy and increase in oxidative stress with the GSC-niche. Thus, it is possible that decreased autophagy flux in GSC may cause their loss due to aging. To investigate this further, we designed an experiment to reduce autophagy levels within germline cells including GSCs, we expressed a double-stranded inverse-repeat (IR) construct designed to target and knockdown (KD) Atg8a (mediated by *Atg8a RNAi*) in the GSCs using *nanosGal4:VP16* driver line (Perkins et al., 2015; Rørth, 1998). GSCs with Atg8aKD were evaluated for oxidative stress, GSC loss, self-renewal capacity and niche occupancy.

We confirmed the depletion of *Atg8a* message using qPCR (**Figure 2A**). In addition, we had previously reported reduced autophagosome formation upon depletion of Atg8a in the GCs (Nilangekar et al., 2019). Autophagy is induced in response to disruption of redox homeostasis to protect cells from apoptosis while impairment of autophagy leads to accumulation of oxidative stress. Hence, we decided to investigate if KD of *Atg8aRNAi* in GCs lead to increased oxidative stress. To test this, we stained *Atg8aRNAi* expressing ovarioles with dihydroethidium (DHE). DHE exhibits blue fluorescence in the cytosol and when oxidized by superoxide molecules intercalates cell’s DNA and stains the nucleus leading to emission of bright red fluorescence at emission maxima 610 nm when excited at 535 nm ((Robinson et al., 1994; Rothe and Valet, 1990). GCs expressing *Atg8aRNAi* within the germaria and developing egg chambers had significantly higher levels of ROS as seen by increase in the red fluorescence as compared to the control (**Figure. 2B-B**”). This increase in red fluorescence was also detected in nurse cells (GCSs) of older egg chambers indicating an increase in the production of superoxide molecules within the cytosol of *Atg8aRNAi* expressing cells. These data suggest that disruption of autophagy leads to oxidative stress within the GCs.

**Figure 2.**
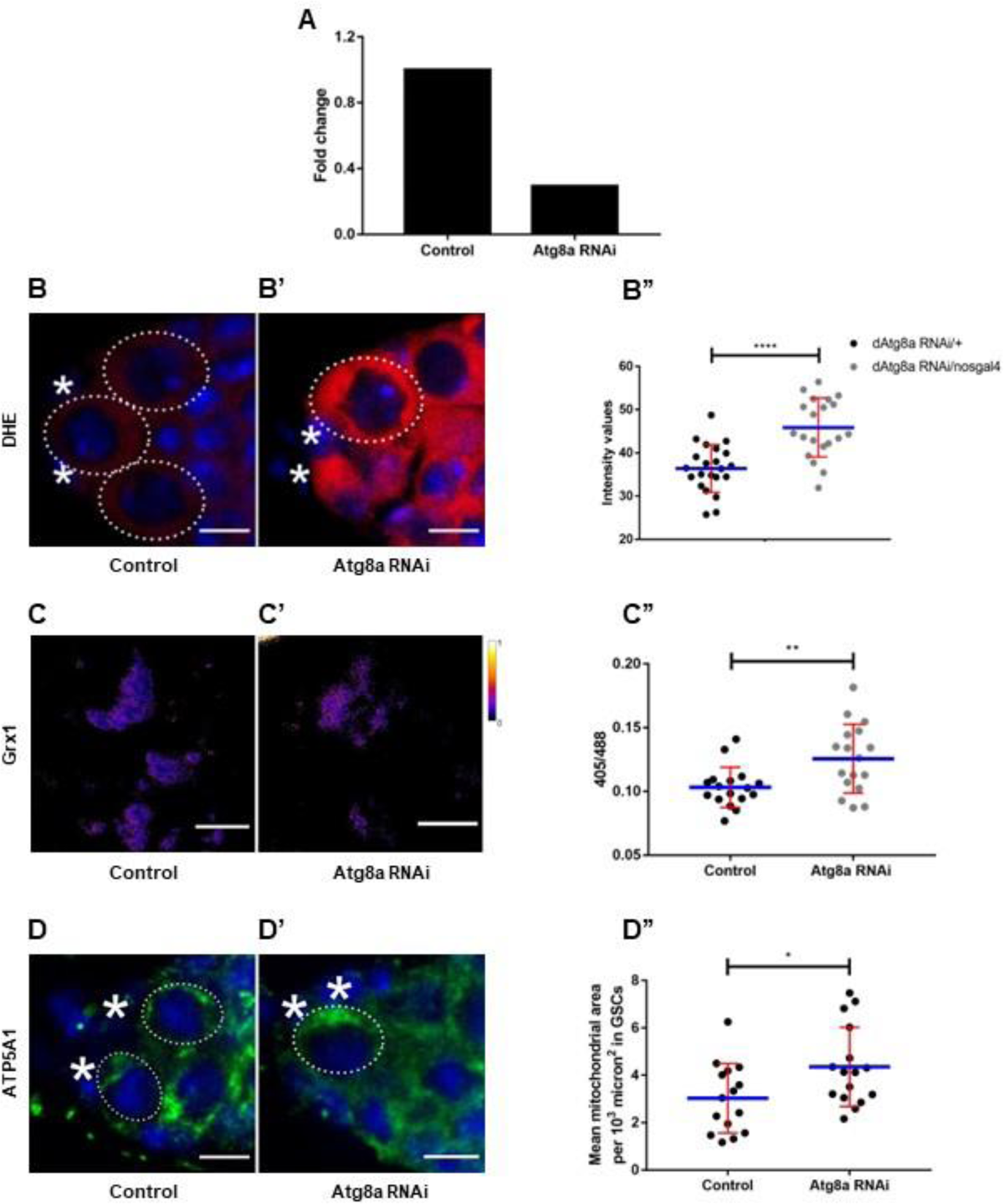
Loss of Atg8a increases reactive oxygen species (ROS) levels. (A)A qPCR analysis shows 70 percent decrease in Atg8a expression in Atg8a RNAi as compared to control. (B-B’’) Knockdown of Atg8a shows elevated ROS levels in the GSCs measured by Dihydroethidium (DHE) (red) Scale bar-5µm. (C-C’) GSCs of indicated genotypes showing ratiometric image of emission of roGFP2. Scale bar-5µm. Knockdown of Atg8a in the GSCs show increase in ROS measured by the change in ratiometric shift in excitation of the redox probe mito-roGFP2 from 488 to 405 (C’’) (n=20). (D-D’’) GSCs of indicated genotypes stained with anti-ATP5α (green) to mark the mitochondria. Mean mitochondrial area increases upon Atg8a knockdown. Error bars represent standard deviation (S.D.) in red and blue lines represent mean, n=20. Control: ywhsFLP1/w; Atg8a RNAi/+; Atg8a RNAi/+, Atg8a RNAi: ywhsFLP1/ywhsFLP12; Atg8a RNAi/+; Atg8a RNAi/nos-Gal4. *p < 0.05, **p < 0.01, ****p < 0.0001

Autophagy is not only important for removal of toxic proteins and protein aggregates, it is also crucial for maintaining mitochondrial homeostasis by removal of non-functional and damaged mitochondria by the process of mitophagy (Bakula and Scheibye-Knudsen, 2020; Lee et al., 2018; Onishi et al., 2021). Damaged mitochondria if not cleared from within the cells generate excess reactive oxygen species (mROS) which can trigger cell death (Shadel and Horvath, 2015). Importantly, as cells, tissues and organisms age there is increased production of mROS due to inefficient clearing of damaged mitochondria by mitophagy. To test if autophagy block leads to mROS accumulation in GSCs, we used a germline specific mitochondrial redox sensor, mito-roGFP2-Grx1, previously characterized by our lab (Albrecht et al., 2014; Nilangekar and Shravage, 2019). This reporter senses the redox state of glutaredoxin (Grx1) within the cells and the redox potential is determined by the 405/488 ratiometric(Pan et al., 2007; Zhao et al., 2008) shift in fluorescence of roGFP2. Ratiometric shift in control and *Atg8aRNAi* expressing GSCs was acquired and quantified. Our analysis shows that the redox state of GSCs with Atg8aKD is significantly higher as compared to the controls (**Figure 2C-C**”). This is indicative of accumulation of damaged and nonfunctional mitochondria in the Atg8a GSCs and germ cells. To test this further, we utilized anti-ATP5alpha (ATP5A1) that marks complex V of the mitochondrial electron transport chain and measured average mitochondrial area in control and *Atg8aRNAi* germaria. In Atg8aKD the average area of mitochondrion was significantly larger as compared to the controls suggesting a disruption in mitochondrial clearance leading to their accumulation (**Figure 2D-D”**). To summarize, these findings indicate that Atg8a KD in GCs including GSCs leads to increased generation of mitochondrial ROS.

Next, we performed GSC retention assay to assess the effect of autophagy disruption. In wild-type *Drosophila* GSCs are turned-over gradually due to aging ((Pan et al., 2007; Zhao et al., 2008)). To test this, we quantified the GSC number at 1,4 and 8 weeks old germaria in both control and Atg8aKD. GSCs in *Drosophila* possess a spherical structure rich in cytoskeletal protein called the “sprectrosome”. Sprectrosome was labelled with anti-alpha-Spectrin antibody and GSCs were imaged using confocal laser scanning microscopy. Atg8aKD in the GSCs led to decrease in the average GSCs within the germarium at 1-week and 4-weeks of age as compared to the controls (**Figure 3A, B**). This indicated that disruption of autophagy led to their accelerated loss from the GSC niche. Interestingly, at 8-weeks of age, a significantly greater number of GSCs were retained in the GSC-niche as compared to controls (**Figure 3A, B**) indicating that autophagy maybe necessary for GSC turnover in old female *Drosophila*. The GSC distribution data also reflected similar findings. GSC niches in *Atg8aRNAi* possessed fewer GSC at 1 and 4-weeks of age as compared to the controls. However, at 8-weeks of age more than 90% of *Atg8aRNAi* GSC-niches possessed one or more GSCs as compared to control wherein the proportion of niches devoid of GSCs was higher (**Figure 3C**). Taken together, our data supports the fact that reduced autophagy accelerates GSC loss from the niche.

**Figure 3:**
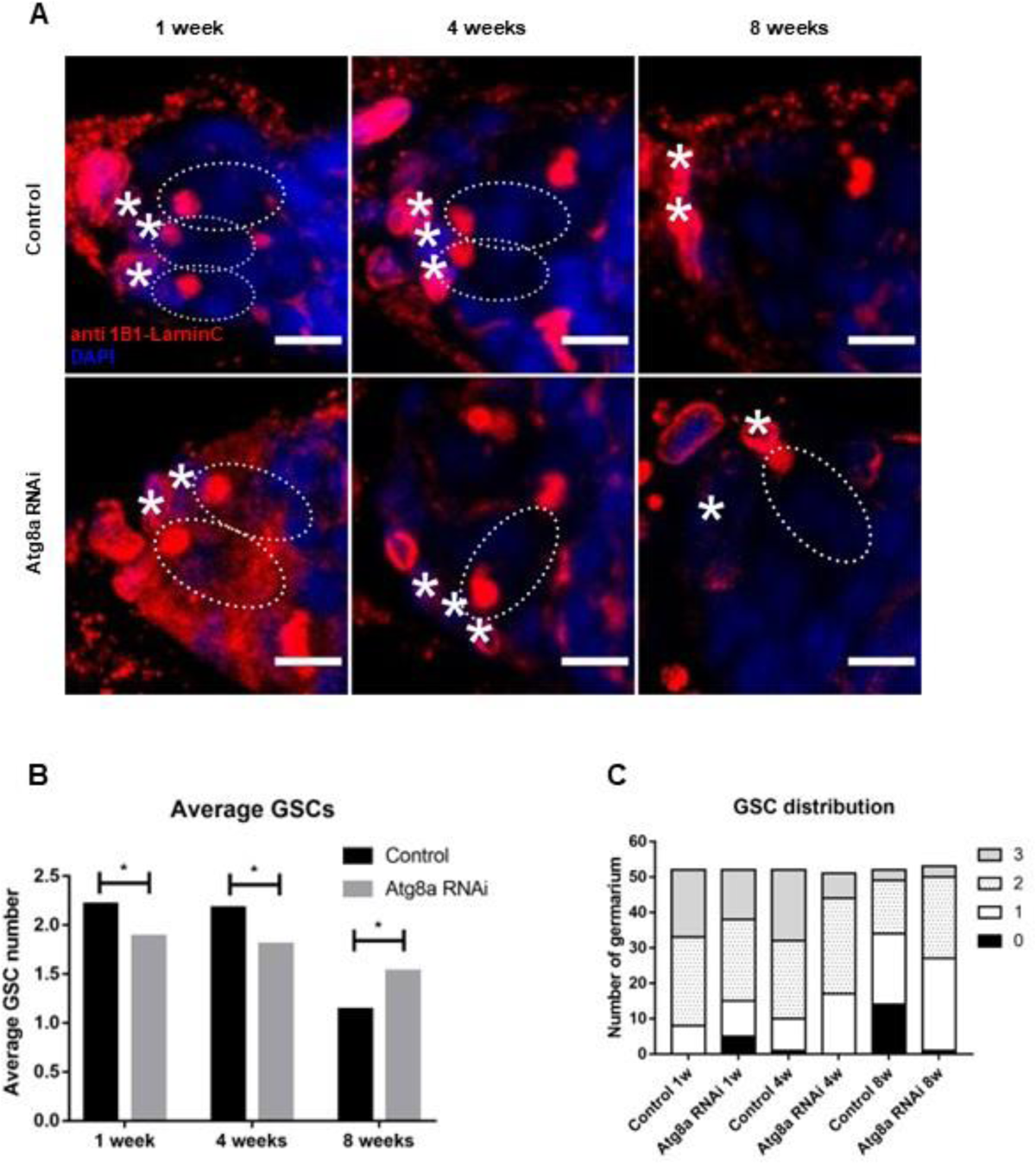
Knockdown of Atg8a in the GSCs lead to decline in GSC counts at 4 weeks. (A) Anterior tip of germarium showing GSCs. Spectrosomes marked by 1B1 (red), cap cells marked by LaminC (red) in controls (Atg8a RNAi/+) (top) and test (Atg8a RNAi/nos-Gal4) (bottom) at 1-8 weeks. Nuclei are marked in blue. Scale bar-5µm. Dotted ovals mark the GSCs and asterisk mark the cap cells. Average GSCs (B) decrease upto 4 weeks and increase at 8 weeks in Atg8a RNAi as compared to respective controls. Stacked bar graph (C) shows distribution of GSCs in germaria of indicated genotypes at 1-8 weeks. Control: ywhsFLP1/w; Atg8a RNAi/+; Atg8a RNAi/+, Atg8a RNAi: ywhsFLP1/ywhsFLP12; Atg8a RNAi/+; Atg8a RNAi/nos-Gal4. n=50, *p < 0.05.

Autophagy block has been shown to influence cell cycle progression and induce replicative stem cell senescence in the *Drosophila* intestine during aging (Laura Garcia Prat 2016, Nagy et al. 2018). We examined if mitosis was affected in GSCs with Atg8a KD. To study GSC division we followed the spectrosome morphology to identify mitotically dividing cells. Spectrosomes go through change in morphology during different phases of cell cycle within the GSCs. For instance, during S phase, GSCs display a “plug,” “elongated,” or “bar” spectrosome morphology. The plug shaped spectrosome is assembled from the newly formed cystoblast and then fused to the original spectrosome thereby connecting the GSC and the cystoblast. During early G2, GSCs exhibit “exclamation point” spectrosome morphology, as the connection between GSCs and the cystoblast is severed. During late G2 and M, GSC fusomes display a “round” fusome (de Cuevas and Spradling, 1998; Kao et al., 2015 and Figure 4A). We counted the number of mitotically active cells characterized by the presence of elongated spectrosomes in Atg8aKD. The number of GSCs undergoing mitosis did not change significantly at 1-week and 4-weeks in Atg8aKD GSCs vs control GSCs. However, at 8 weeks of age, GSCs with Atg8aKD exhibited a higher mitotic index as compared to the controls (**Figure 4A, B-B”**).

**Figure 4:**
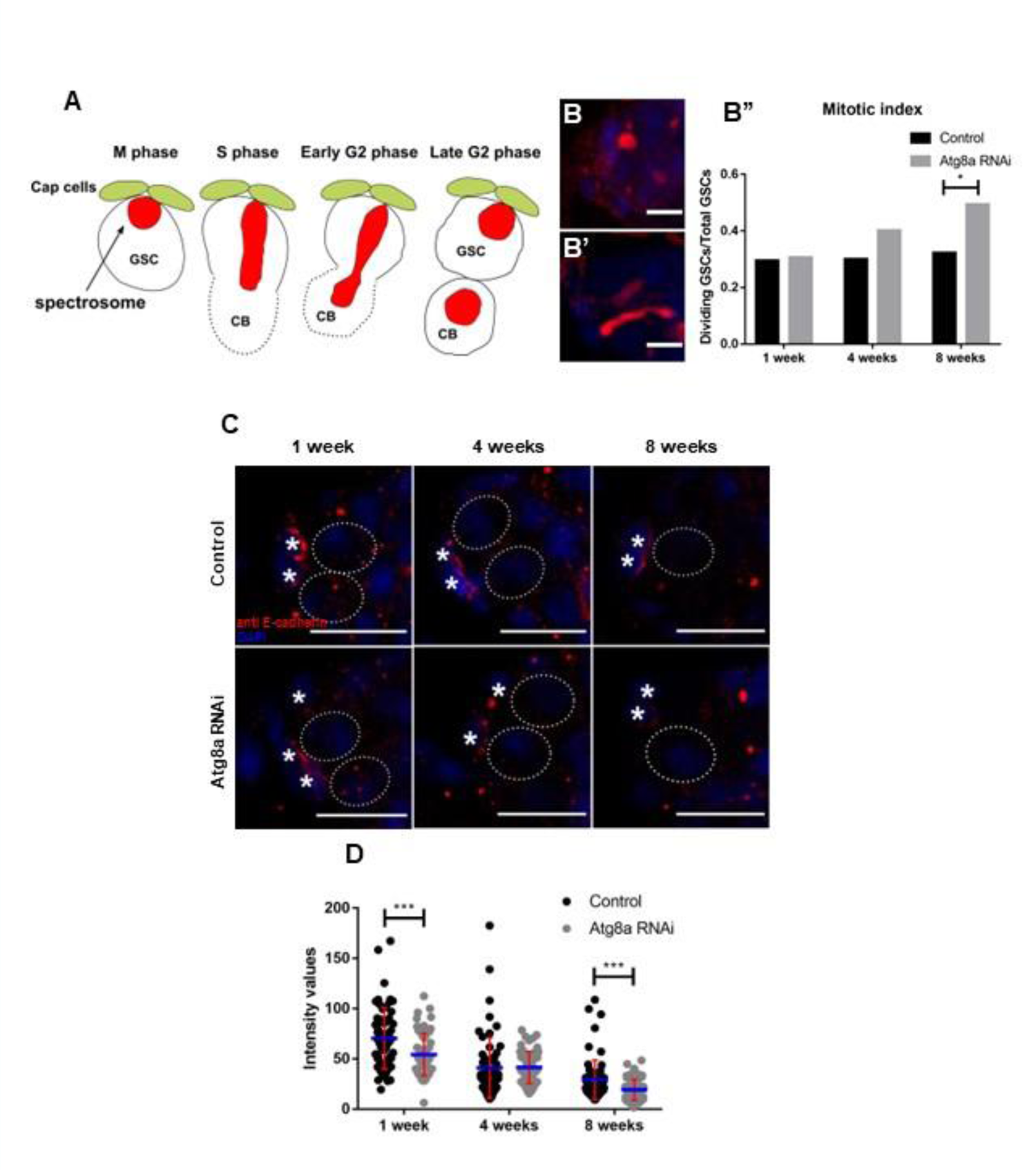
Effect of Atg8a knockdown in GSC-niche occupancy and GSC activity. (A) Schematic representation of shape of spectrosome during different stages of GSC division. (B-B’’) Spectrosomes (red) after mitosis in GSCs (B) and elongated spectrosomes in dividing GSCs (B’) Scale bar-5µm. GSC activity measured by mitotic index increases significantly at 8 weeks in the Atg8a RNAi (B’’). (C) Reduced E-cadherin (red) at GSC-niche in controls (UAS Atg8a RNAi/+) (top) and test (UAS Atg8a RNAi/nos-Gal4) (bottom) at 1-8 weeks. Nuclei are marked in blue. Dotted ovals mark the GSCs and asterisk mark the cap cells. Scale bar-10µm. (D) Decreased E-cadherin intensity upon Atg8a knockdown as compared to controls at 1 and 8 weeks. Error bars represent standard deviation (S.D.) in red and blue lines represent mean, n=20. Control: ywhsFLP1/w; Atg8a RNAi/+; Atg8a RNAi/+ Atg8a RNAi: ywhsFLP1/ywhsFLP12; Atg8a RNAi/+; Atg8a RNAi/nos-Gal4. *p < 0.05, ***p < 0.001

GSCs are maintained by extrinsic signaling from the niche cells in particular the terminal filament (TFs) cells and the cap cells (CCs). Dpp (BMP2/4 homolog in *Drosophila*) is secreted from the TFs and CCs and is essential for GSC self-renewal and controls GSC divisions (Xie and Spradling, 1998). Dpp signal is further transduced into the GSCs via phosphorylation of Mad (phospho-Smad or pSmad) in *Drosophila* (Pan et al., 2007; Raftery and Sutherland, 1999). Previous studies have reported a decline in phosphorylation of Mad in aged GSCs, indicative of impaired Dpp signaling activity (Pan et al., 2007). Reduced Dpp signaling activity contributes to GSCs loss due to aging (Pan et al., 2007; Zhao et al., 2008). We examined if the phosphorylation of Mad is regulated by Atg8a. In *Atg8aRNAi* the pMad levels were unaltered between the control and test GSCs (**Supplementary Figure 2)**. However, at 8 weeks of age, we detected a significant reduction of staining intensity for pMad. These data suggest that Atg8a does not influence phosphorylation of Mad until 4-weeks of age and GSC loss in *Atg8aRNdi* is independent of pMad signaling.

Cell adhesion mediated by E-cadherin, (DE-cadherin in *Drosophila*) encoded by the *shotgun (shg)*, plays an important role during oogenesis and mediates cell-adhesion for anchoring GSCs to cap cells within the GC-niche (Song et al., 2002). The accumulation of E-cadherin between GSC-niche boundaries declines in an age-dependent manner and is partly responsible for age-dependent loss of GSCs (Pan et al., 2007; Tseng et al., 2014). In *Atg8aKD* there is a significant decrease in E-cadherin at the cap cell-GSC contact sites as compared to controls (**Figure 4C, D**). At 4 weeks of age, the decrease in E-cadherin levels was not significant amongst the control and *Atg8aRNAi* expressing GSC-niche contact sites. However, E-cadherin levels at 8 weeks of age continued to decrease further and even though there are higher numbers of GSCs in *Atg8aRNAi* as compared to the controls (**Figure 4D**). To summarize, E-cadherin expression is reduced at GSC-cap cell contact sites upon Atg8aKD in GSCs indicating that Atg8a influence their niche occupancy.

### Atg8a ^OE^ specifically in the germarium leads to increase in autophagic flux and reduced generation of mitochondrial ROS

Previous studies in various model systems including *Drosophila* suggest that upregulating autophagy flux prolongs life span (Aparicio et al., 2019; Bjedov et al., 2020; Simonsen et al., 2008). These studies showed that increased autophagy reduced the accumulation of toxic entities within the cytoplasm including protein aggregates and dysfunctional organelles, and promoted cell health. Studies in yeast have shown that Atg8 overexpression leads to increase autophagosome size and activity (Nair et al., 2012). Our hypothesis is that increasing autophagy flux specifically within the GSCs will prevent their loss from the GSC-niche by maintaining cellular homeostasis. In order to delay the loss of GSCs due to aging we decided to overexpress *Atg8a* specifically within the germline cells including GSCs. To achieve overexpression of *Atg8a* within the GSCs, *mCherry-Atg8a* expression was driven by *nanos-VP16Gal4* (*UASp-mcherry-Atg8a X nanosVP16-Gal4*) (Rørth, 1998). The first step was to determine if *Atg8a^OE^* increases autophagy flux within the germline cells. Since autophagy is a multi-step process, the use of autophagosome-lysosome fusion inhibitors makes it possible to study the dynamic changes in the autophagic process. In particular it is possible to quantify the increase in autophagosome formation accompanied by an increase in the number of lysosomes which together leads to increased formation of autolysosomes (Klionsky et al., 2021). Investigation of autophagic flux was carried out in the presence of autophagosome-lysosome fusion inhibitor, chloroquine (CQ).

We confirmed *Atg8a^OE^* using qPCR. The *Atg8a* mRNA was upregulated 7-fold in *Atg8a^OE^* as compared to controls (**Supplementary Figure 3A**). In addition, we monitored the expression of mCherry within the GCs as Atg8a is tagged with mCherry at its N-terminal. As compared to the control the mCherry fluorescence was found to be increased 10-fold (**Supplementary Figure 3B**). Ovaries were dissected from *Atg8a^OE^* flies and subjected to autophagy flux (described in materials and methods). Anti-CathepsinL staining was used to mark lysosomes in the *Atg8a^OE^* flies. mCherry-Atg8a puncta [(mCherry-CathepsinL)-CathepsinL] and mCherry-CathepsinL colocalized puncta were considered as a read-out of the number of autophagosomes and autolysosomes respectively, in the presence and absence of chloroquine (CQ) treatment. The number of autolysosomes increase significantly in the germarium but not GSCs expressing Atg8a in untreated conditions (**Figure 5A-C**). We observed inhibition of fusion between autophagosomes and lysosomes following CQ treatment in *Atg8a^OE^*. This resulted in accumulation of mCherry-CathepsinL colocalized puncta in both control and *Atg8a^OE^* as compared to the fed controls (**Figure 5A-C)**. The differences in autolysosomes in GSCs in untreated and CQ treated germaria were not significant, however, an increase in autophagy flux could be detected. An autophagosome and lysosome fuse to form a single vesicle, called the autolysosome, during the final stages of the autophagic process (**Figure 5D, E**). In absence of CQ, *Atg8a^OE^* led to formation of large sized autophagosomes and lysosomes (**Figure 5B, D** and **Supplementary Figure 3E**). In presence of CQ, we observed the accumulation of unfused mCherry-Atg8a decorated by CathepsinL positive vesicles in *Atg8a^OE^* germaria (**Figure 5E**). To get further insights, a size cut-off for autolysosome was assigned in order to compare the results between treated and untreated samples. Autolysosomes and lysosomes were grouped into two groups based on their area, smaller sized vesicles with 0.03-1um^2^ area and large sized vesicles with 1-3um^2^ (**Supplementary Figure 3C-K**). *Atg8a^OE^* germaria showed an increase in number and size of mCherry-Atg8a vesicles (autophagosomes) and CathepsinL puncta upon CQ treatment (**Figure 5A-C** and **Supplementary Figure 3C-E and 3G-I**). However, in *Atg8a^OE^* GSCs, we observed no significant changes in the number of smaller sized mCherry-Atg8a puncta, and formation of larger sized mCherry-Atg8a puncta in GSCs did not occur (**Supplementary figure 3F**). In GSCs, smaller sized CathepsinL puncta increased in both fed and CQ conditions (**Supplementary figure 3G, J**), however, no significant change in large sized CathepsinL puncta in GSCs was observed (**Supplementary Figure 3K**). This analysis suggest that autophagy flux increased in the germline cells including GSCs upon *Atg8a^OE^*.

**Figure 5:**
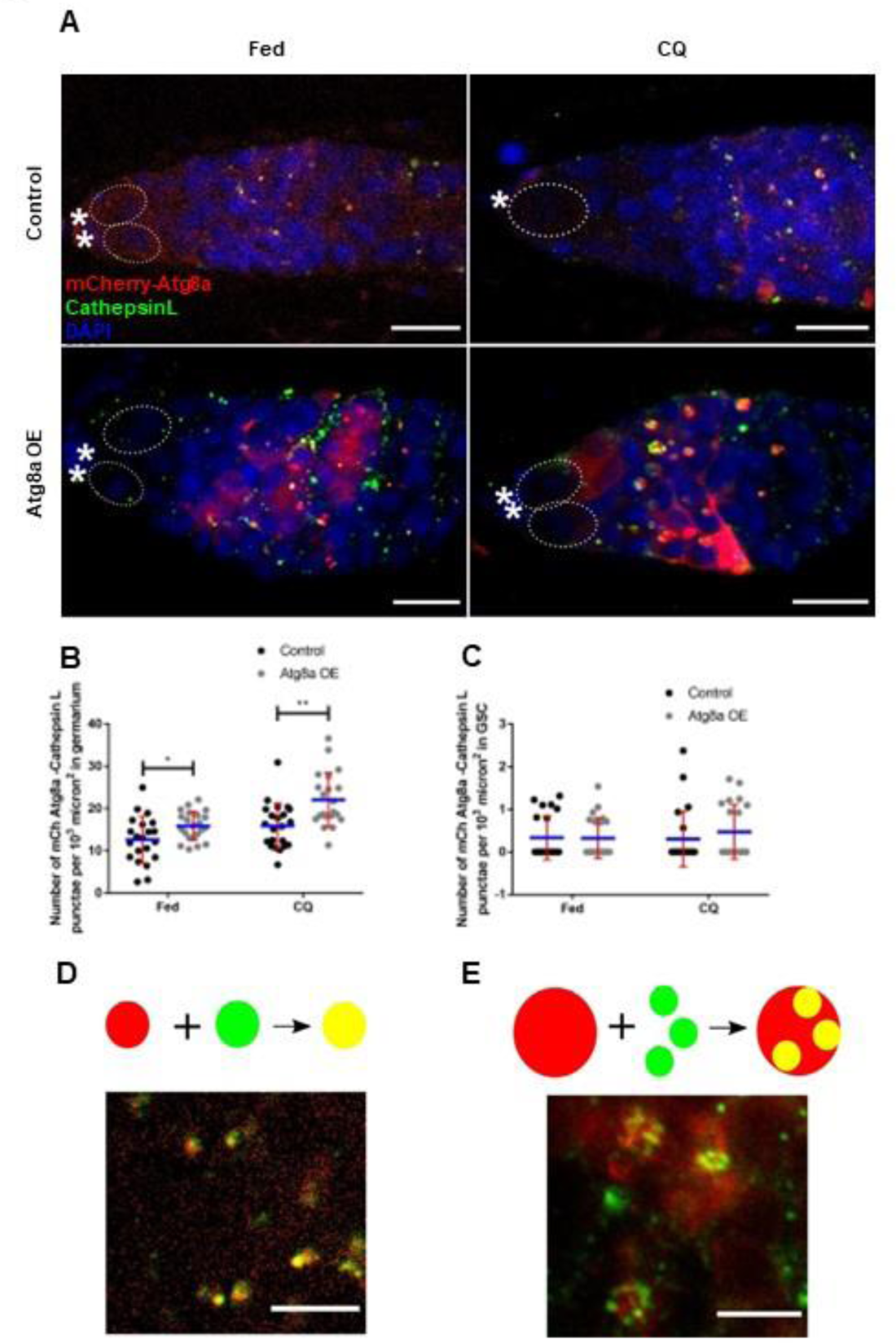
Autophagic flux increases upon Atg8a overexpression in the GSCs. (A) Representative images for indicated genotype fed on yeast and chloroquine (left to right) stained for CathepsinL depicting colocalization between mCherry-Atg8a-CathepsinL to mark the autolysosomes (yellow). Nuclei are marked in blue. Dotted ovals mark the GSCs and asterisk mark the cap cells. Scale bar-10µm. (B-C) Increase in mCherry Atg8a-CathepsinL colocalization in germarium (B) and GSCs (C). Error bars represent standard deviation (S.D.) in red and blue lines represent mean. n=20. (D-E) Fusion between autophagosomes and lysosomes to form a single vesicle (yellow) (D) and inhibition of fusion between autophagosomes and lysosomes upon CQ treatment (E). Scale bar − 5 µm. Control: yw;UASp mCherry Atg8a/nosmCherry Atg8a;+, Atg8a OE: yw/w;UASp mCherry Atg8a/nosmCherry Atg8a; nos-Gal4/+. *p < 0.05, **p < 0.01

To test if autophagy block leads to mROS accumulation in GSCs, we used a germline specific mitochondrial redox sensor, mito-roGFP2-Grx1, previously characterized by our lab (Albrecht et al., 2014; Nilangekar and Shravage, 2019). Our analysis shows that the redox state of GSCs with *Atg8aOE* is significantly lower as compared to the controls (**Figure 6A-A**”). Previous studies have shown that promoting mitochondrial fission or elevated autophagy leads to reduction in size of mitochondria which facilitates mitophagy (Bjedov et al., 2020; Rana et al., 2017). We investigated if mitochondrial size is reduced in *Atg8a^OE^* mitochondrial area in control and *Atg8a^OE^* by using anti-ATP5A1 antibody to visualize mitochondria. The average area of mitochondria was much smaller in *Atg8a^OE^* as compared to controls (**Figure 6B-B”**). Smaller mitochondrial size facilitates mitochondrial clearance and this may be the case in *Atg8a^OE^*. Thus, *Atg8a^OE^* could facilitate removal of damaged mitochondria by regulating their size thereby maintaining redox homeostasis. Taken together these findings suggest that *Atg8a^OE^* upregulates autophagy and facilitates redox homeostasis during aging.

**Figure 6:**
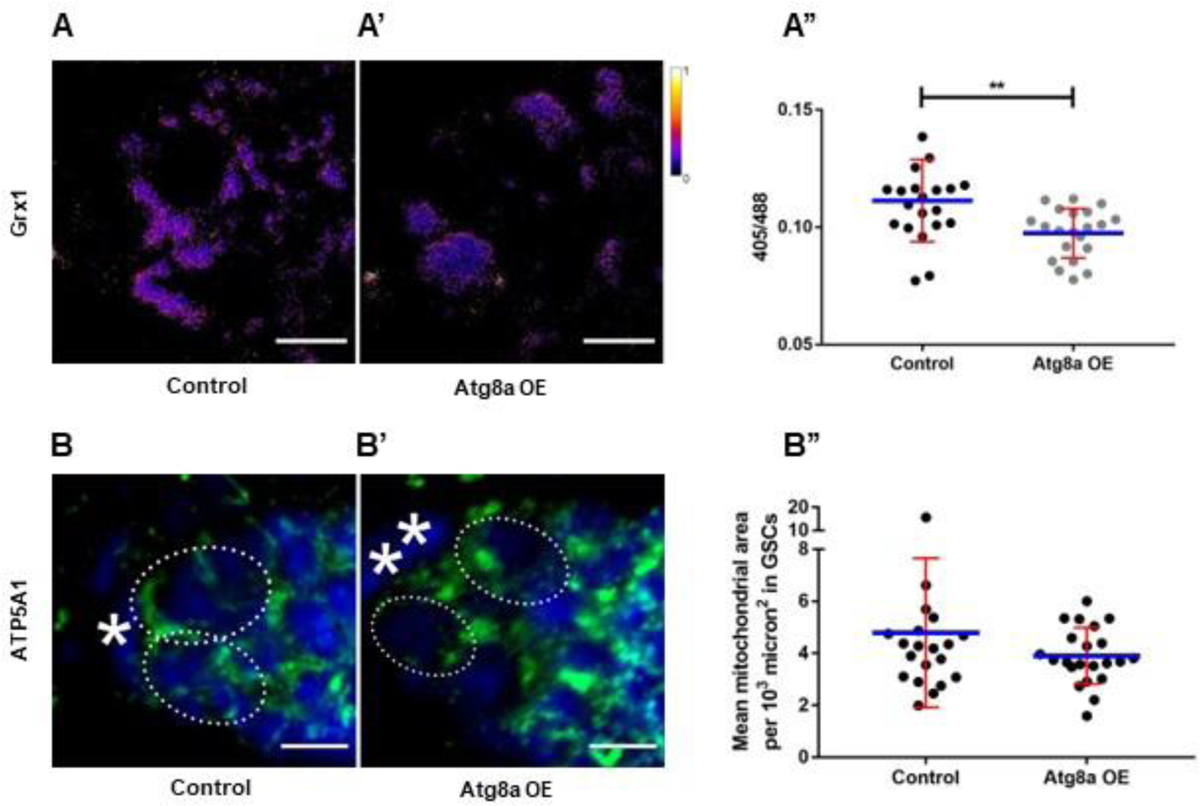
Atg8a overexpression affects mitochondria in GSCs (A-A’’) Overexpression of Atg8a in the GSCs show decrease in ROS measured by reduction in 405/488 ratio (A’’) (n=20). Scale bar-5µm. GSCs of indicated genotypes stained with anti-ATP5α (green) to mark the mitochondria (B-B’). Mean mitochondrial area reduces upon Atg8a overexpression in the GSCs as compared to controls (B’’). Scale bar-5µm.Error bars represent standard deviation (S.D.) in red and blue lines represent mean. n=20. (A-A’’) Control: yw; UASp mCherry Atg8a/nosP-mito-roGFP2-Grx1;+, Atg8a OE: yw/w; UASp mCherry Atg8a/ nosP-mito-roGFP2-Grx1; nos-Gal4/+. (A-B’’) Control: yw; UASp mCherry Atg8a/+, Atg8a OE: yw/w; UASp mCherry Atg8a/+; nos-Gal4/+. **p < 0.01

### Atg8a^OE^ leads to longer retention of GSCs in middle-aged flies due to increased expression of E-cadherin in the GSC-niche

Our data show that increasing autophagy specifically within the GSCs increased autophagic flux. Previous studies demonstrated that elevated autophagy levels are beneficial and promote cell health and longevity (Aparicio et al., 2019; Bjedov et al., 2020; Simonsen et al., 2008). However, very high levels of autophagy are detrimental to the cells and can lead to cell death. We tested if increased autophagy levels in *Atg8a^OE^* are beneficial for GSCs. In particular, if *Atg8a^OE^* in GSCs slows their decline in number due to aging and exhibits prolonged mitotic activity. To test this, we quantified the GSC number at 1,4 and 8-weeks old germaria in both control and *Atg8a^OE^*. We observed a significant increase in average GSC counts in 1-week and 4-weeks old *Atg8a^OE^* followed by sharp decline at 8-week (**Figure 7A, B**). To get a deeper insight, GSC numbers within every GSC-niche were counted. **Figure 7C** shows the GSC distribution in control and *Atg8a^OE^*. At 1-week of age there is a significant increase in the number of germaria carrying three GSCs in *Atg8a^OE^* as compared to control. A few germaria possessing four GSCs were also observed (**Figure 7C**). At 4 weeks of age, in control the proportion of germaria with one GSC was more due to loss of GSCs upon aging. In contrast, *Atg8a^OE^* exhibited a higher number of germaria with two or more GSCs at 4-weeks of age. This situation was completely reversed at the 8-weeks’ time point. In controls at 8-weeks the loss of GSCs was gradual as reported for wild type previously (Pan et al., 2007). Unexpectedly, at 8-weeks in Atg8a^OE^ there was a significant increase in GSC loss and many of the germaria did not possess even a single GSC. This indicates that elevated levels of autophagy in aged germaria are detrimental to the GSCs. Taken together these data suggest that elevated levels during early and mid-age have positive influence on the GSCs while in old age higher levels of autophagy promote GSC loss.

**Figure 7:**
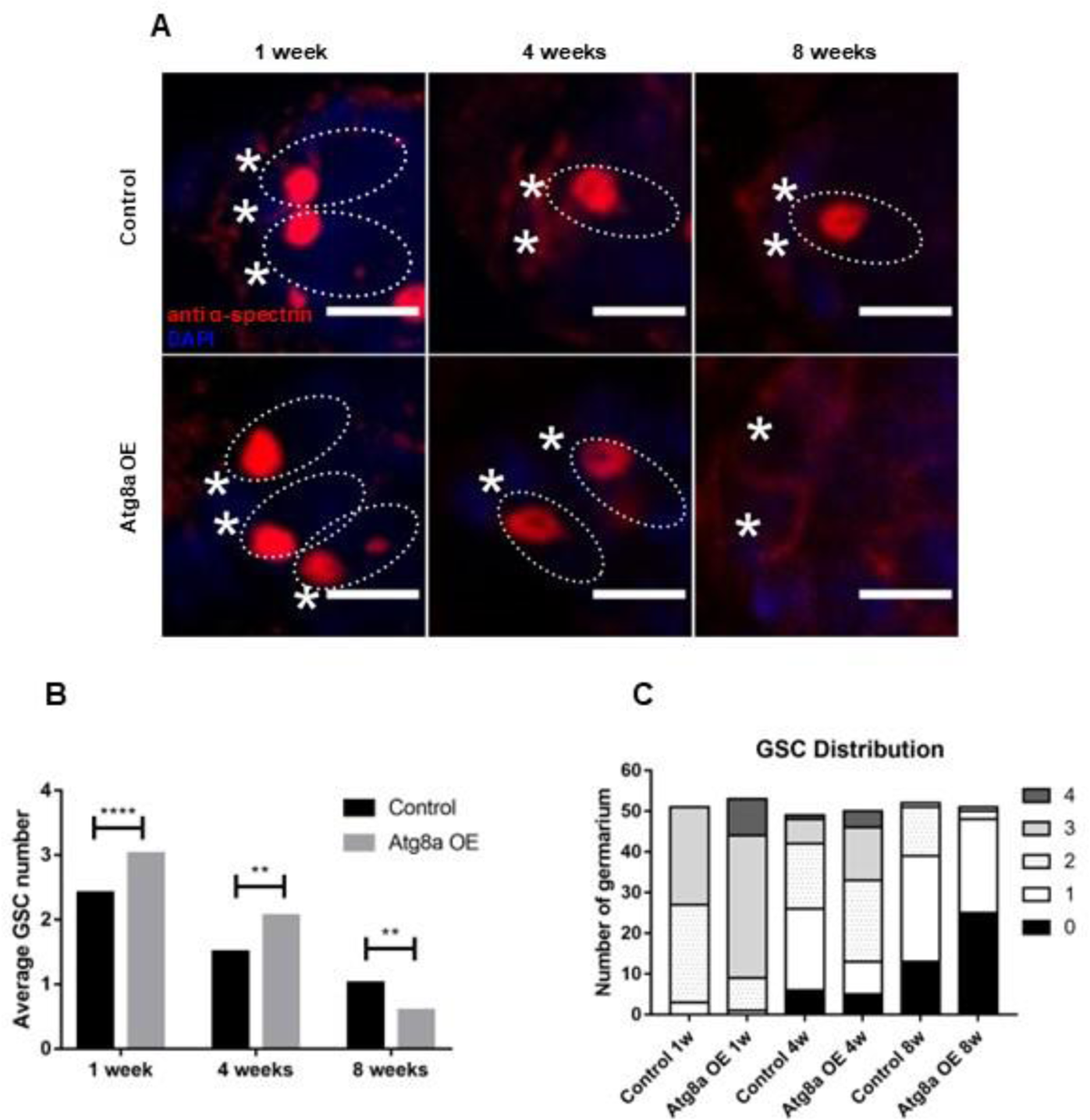
Overexpression of Atg8a in the GSCs increase GSC counts upto 4 weeks. (A) Spectrosomes marked by alpha spectrin (red) in controls (UASp mChAtg8a/+) (top) and test (UASp mChAtg8a/nos-Gal4) (bottom) at 1-8 weeks. Nuclei are marked in blue. Dotted ovals mark the GSCs and asterisk mark the cap cells. Scale bar-5µm. (B-C) Average GSCs (B) increase upto 4 weeks and decrease at 8 weeks in Atg8a overexpression as compared to respective controls. Stacked bar graph (C) shows distribution of GSCs in germaria of indicated genotypes at 1-8 weeks, n=50. Control: yw; UASp mCherry Atg8a/+, Atg8a OE: yw/w; UASp mCherry Atg8a/+; nos-Gal4/+**p < 0.01, ****p < 0.0001

Accelerated GSC loss at 8-weeks could be due to increased differentiation or induction of GSC death. *Atg8a^OE^* at 1-week and 4-week does not lead to accelerated differentiation of GSCs so this possibility could be ruled out. GSCs in *Drosophila* rarely undergo cell death as upto 50% of GSCs remain in the germarium at 8-weeks of age (Pan et al., 2007). However, certain genetic mutations can induce programmed cell death in GSCs (Kober et al., 2019). Upon induction of PCD GCs were shown to be positive for cleaved caspase-3(Peterson et al., 2003). We stained germaria with cleaved caspase-3 at 1, 4 and 8-week age. No significant differences were detected in cleaved caspase-3 staining at 1 and 4-week old control and *Atg8a^OE^* germaria (**Supplementary Figure 4A, A’**). However, a significant increase in Caspase 3 positivity was detected in Atg8a^OE^ GSCs and germaria at 8 weeks as compared to controls (**Supplementary Figure 4A, A”)**. This indicates that in aged germaria *Atg8a^OE^* promotes programmed cell death in GSCs and germaria.

We counted the number of mitotically active cells characterized by the presence of elongated spectrosomes (**Figure 8A**). Our analysis showed elevated mitotic activity significantly at all three 1week, 4 week and 8-week time points tested (**Figure 8A-A”)**. The increase in GSC division was significant in middle aged flies (4 weeks old) in *Atg8a^OE^* (**Figure 8A’’**). Interestingly, even though there are fewer GSCs remaining in the niche at 8-weeks, these GSCs exhibited higher mitotic index as compared to the controls of the same age. These observations suggest that elevated autophagy promotes GSC cell division throughout the life span of the female flies. Our data so far suggests *Atg8a^OE^* within the GSCs led to their longer retention and exhibited increased mitotic activity (**Figure 8A, B**).

**Figure 8:**
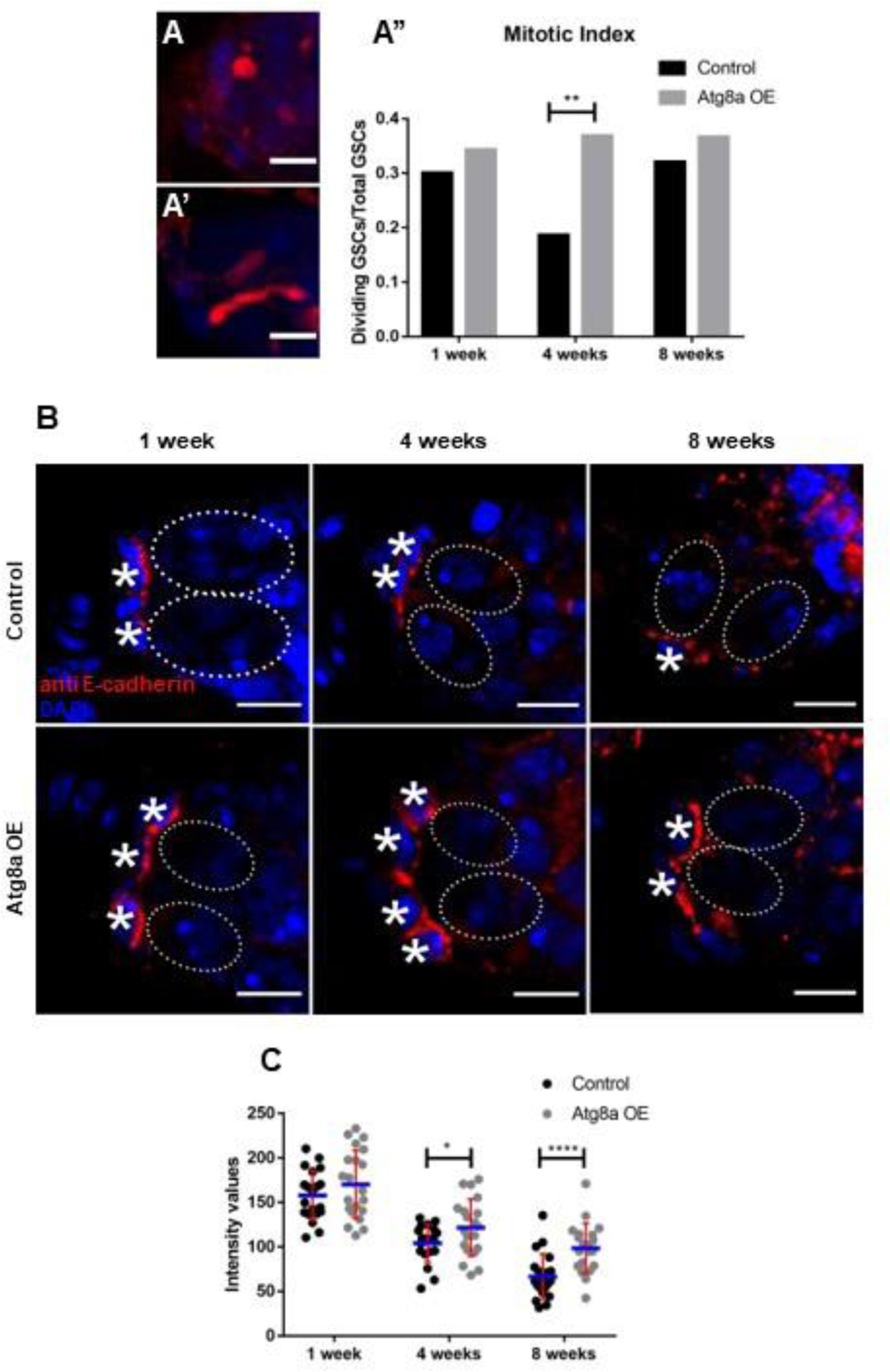
Atg8a overexpression increases GSC activity and E-cadherin at GSC-niche (A-A’’) Spectrosomes (red) after mitosis in GSCs (A) and elongated spectrosomes in dividing GSCs (A’) Scale bar-5µm. Mitotic index of GSCs increases significantly at 4 weeks in Atg8a overexpression (A’’). n=50. (B) E-cadherin (red) at GSC-niche (red) in controls (UASp mChAtg8a/+) (top) and test (UASp mChAtg8a/nos-Gal4) (bottom) at 1-8 weeks. Nuclei are marked in blue. Dotted ovals mark the GSCs and asterisk mark the cap cells. Scale bar-5µm. (B-C) Increase in E-cadherin intensities upon overexpression of Atg8a. Error bars represent standard deviation (S.D.) in red and blue lines represent mean. n=20. Control: yw; UASp mCherry Atg8a/+, Atg8a OE: yw/w; UASp mCherry Atg8a/+; nos-Gal4/+ *p < 0.05, **p < 0.01, ****p < 0.0001

*Atg8a^OE^* leads to a significant increase in pMad levels as compared to controls at 1-week of age. At 4-weeks of age, the pMad levels showed a decrease, however, the differences between control and *Atg8a^OE^* are not significant (**Supplementary Figure 5A-A”**). In contrast, at 8-weeks of age we observed a significant decline in pMad levels as compared to controls. (**Supplementary Figure 5A”)**. These data suggest that there is no direct correlation between *Atg8a^OE^* and phosphorylation of Mad. These data suggest that GSC retention and loss in *Atg8a^OE^* respectively occurs independent of pMad signaling. We wanted to investigate if differences in retention of GSCs within the niche in *Atg8a^OE^* was due to changes in E-cadherin levels during the life span of *Drosophila*. At 1-week of age, there is an increase in the levels of E-cadherin at the cap cell-GSC contact site in *Atg8a^OE^* as compared to controls. At 4-weeks of age, we observed a significant increase in E-cadherin levels in *Atg8a^OE^* as compared to controls. Further, E-cadherin levels remained significantly higher at 8-weeks in *Atg8a^OE^* even though there are fewer GSCs in *Atg8a^OE^* as compared to the controls (**Figure 8B-C**). These data suggest that *Atg8a^OE^* directly or indirectly influences E-cadherin expression at the GSC-niche contact sites. Taken together, these data suggest that the autophagy-mediated longevity of GSCs is independent of Dpp signaling but dependent on E-cadherin.

## Discussion

Loss of proteostasis is considered to be one of the major hallmarks of aging including aging of stem cells. Autophagy is significantly reduced in aged cells, tissues and organisms including humans (Aman et al., 2021; López-Otín et al., 2013). However, the underlying mechanisms of decrease in autophagy and how autophagy influences other factors that affect aging of stem cells is not understood. We have used *Drosophila* females germline stem cells as a model to decipher the role of autophagy in stem cells and their aging (Ishibashi et al., 2020). Our lab and others have shown that basal autophagy in GSCs occurs at very low levels (Nilangekar et al., 2019; Zhao et al., 2018). We measured autophagy in aged GSCs and observed that the basal autophagy in GSCs declines further during the life span of the fly (**Figure 1**).

We demonstrate that Atg8a^OE^ in the female germline stem cells promotes their longer retention as well as activity within the niche by elevating autophagy that facilitates bulk clearance of protein aggregates and removal of damaged organelles. Similarly, Atg8a overexpression, Ref(2)P/p62 overexpression, moderate Atg1 overexpression and Atg5 overexpression have been demonstrated to extend life span (Aparicio et al., 2019; Bjedov et al.; Pyo et al., 2013; Simonsen et al., 2008). This extension of life span was shown to be dependent on improved proteostasis. Atg8a^OE^ is associated with an increase in both the number and size of autophagosomes and this is also documented in this study (Jin and Klionsky, 2014; Nair et al., 2012; Xie et al., 2008). Numerous lysosomes were seen associated with the large-sized autophagosomes detected in Atg8a^OE^ GSCs and germaria leading to an increase in the autophagy flux. The larger autophagosomes carry more cargo for destruction. These observations are in congruence with the reported studies wherein proteostasis was maintained in Atg8a OE, Ref(2)P/p62 OE, moderate Atg1 OE, and Atg5 OE. Maintaining protein homeostasis is crucial for cellular metabolic activity and health. In contrast, *Atg8aRNAi* in the germline cells including GSCs leads to reduction in autophagy and this has a negative impact on GSC maintenance (Nilangekar et al., 2019). Fewer GSCs are retained in the niche with age suggesting an important role of Atg8a in GSC aging.

Redox homeostasis is crucial for maintaining cell health. ROS like 0_2_^-^, H_2_O_2_ and OH^-^ are byproducts of metabolism mostly generated from mitochondria and their aberrant and increased production of can lead to detrimental effects (Scherz-Shouval and Elazar, 2011). Reactive oxygen species damage macromolecules like proteins, lipids and nucleic acids and render them inactive leading to cell death. We observed in Atg8a^OE^ the oxidation of glutaredoxin-1 as measured using roGFP2-Grx1 was significantly reduced suggesting that elevated levels of autophagy favor removal of damaged mitochondria from GSCs. In contrast, in *Atg8aRNAi* GSCs exhibited increased oxidation of the roGFP2-Grx1 sensor. Our observations are similar to those observed in Atg8a OE in neurons, Ref(2)P/p62 OE, moderate Atg1 OE, and Atg5 OE where improved mitochondrial health was crucial for the beneficial effect on lifespan extension. Moreover, the improved mitochondrial homeostasis translated into pro-longevity metabolic profile with improved utilization of lipids for generation of ATP (Bjedov et al.; Ulgherait et al., 2014). Although we have not tested these it is tempting to speculate that Atg8aOE in GSCs promotes efficient utilization of lipids. Indeed, research in male germline stem cells in *Drosophila* has demonstrated accumulation of lipids within the cytoplasm when dysfunctional mitochondria accumulate (Amartuvshin et al., 2020; Sênos Demarco et al., 2019; Sênos Demarco et al., 2020). In contrast, *Atg8aRNAi* showed enhanced oxidation of glutaredoxins suggesting defects in mitochondrial health. Thus, Atg8a^OE^ is beneficial to the GSCs and promotes their health by maintaining proteostasis, and mitochondrial and redox homeostasis.

We demonstrate that *Atg8a^OE^* positively regulates GSC maintenance and aging. This is the first such report in *Drosophila*. Previous studies in intestinal stem cells and male GSCs have shown accelerated aging and loss of stem cells in autophagy mutants and RNAi studies (Ames et al., 2017; Nagy et al., 2018; Sênos Demarco et al., 2019; Zhao et al., 2018). The positive influence of *Atg8a^OE^* on GSC maintenance can be observed till 4 weeks of age. Beyond middle age, elevating autophagy levels appears to be detrimental to the GSCs. GSCs rapidly lost from the niche at 8-weeks of age. This has also been observed in *C. elegans* wherein increased autophagy induction in aged animals is harmful (Wilhelm et al., 2017). It is also possible that the levels of autophagy achieved *Atg8a^OE^* in our study are high to moderate and this can induce cell death in aged females. High levels of autophagy induced by Atg1 can lead to cell death in wing disc cells and have been recently shown to reduce life span in *Drosophila* (Bjedov et al., 2020; Scott et al., 2007). Indeed, as reported previously in wing disc cells we found an increase in cleaved caspase-3 activity at 8-weeks of age. These data suggest that timing of expression and the levels of autophagy induction are crucial for the positive influence of autophagy on GSC maintenance and aging. In contrast, *Atg8aRNAi* did not show any significant effect on the GSC maintenance and aging until 4 weeks of age. However, at 8-week higher number of GSCs were detected in the niche as compared to the controls indicating that GSC loss at 8-week could be driven by autophagy. This finding is supported by the fact that more autophagosomes and autolysosomes are observed in GSCs at 8 weeks as compared to 4 weeks of age (**Figure 1**). How autophagy is upregulated and by what mechanism in the aged tissue is an open question.

Previous studies in stem cells have shown that autophagy regulates proliferation of stem cells. Nagy and colleagues have demonstrated that loss of autophagy leads to DNA damage which activates Chk2 Leading to cell cycle arrest and apoptosis (Nagy et al., 2018). *Atg8a^OE^* GSCs had higher mitotic index which was more significant at mid age. Research in Hematopoietic stem cells of murine origin has shown that autophagy is important to maintain regenerative capacity of old HSCs (Ho et al., 2017). However, in *Atg8aRNAi* GSC divisions occurred at a significantly higher rate at 8-week of age which is perplexing. It is possible that both the levels of autophagy and timing of expression of autophagy may have differential effects on GSC division. In addition, GSC heterogeneity may contribute to this phenotype (Ng et al., 2018). Further experimentation is necessary to understand the complexity of the observations made in both *Atg8a^OE^* and *Atg8aRNAi*.

*Atg8a^OE^* GSCs exhibited higher levels of E-cadherin suggesting stronger niche occupancy while Atg8a RNAi expressing GSCs showed reduced E-cadherin expression. This is in contrast to what is known in cancer stem cells where autophagy machinery is involved in degradation of E-cadherin in breast cancer cells(Damiano et al., 2020). However, it has been demonstrated that the tumor suppressor function of Beclin1 (Atg) in breast cancer cells requires E-cadherin (Wijshake et al., 2021). Thus, the regulation of expression of E-cadherin by Atg genes could be context dependent. It is possible that E-cadherin trafficking is influenced by autophagy upregulation in GSCs and this needs further investigation. It has been shown that Rab11 regulates GSC maintenance and it is required for trafficking of E-cadherin from the GSC cytoplasm to the GSC-niche cell contact sites (Bogard et al., 2007). The expression and distribution of Rab11 in *Atg8a^OE^* would be provide crucial knowledge on how E-cadherin trafficking and its recylcing is affected. Taken together, our work supports the observation that controlled upregulation of autophagy is beneficial to the survival and activity of stem cells and use of autophagy modulators would be critical in stem cell regenerative therapies for better outcomes.

---

**Supplementary figure 1:**
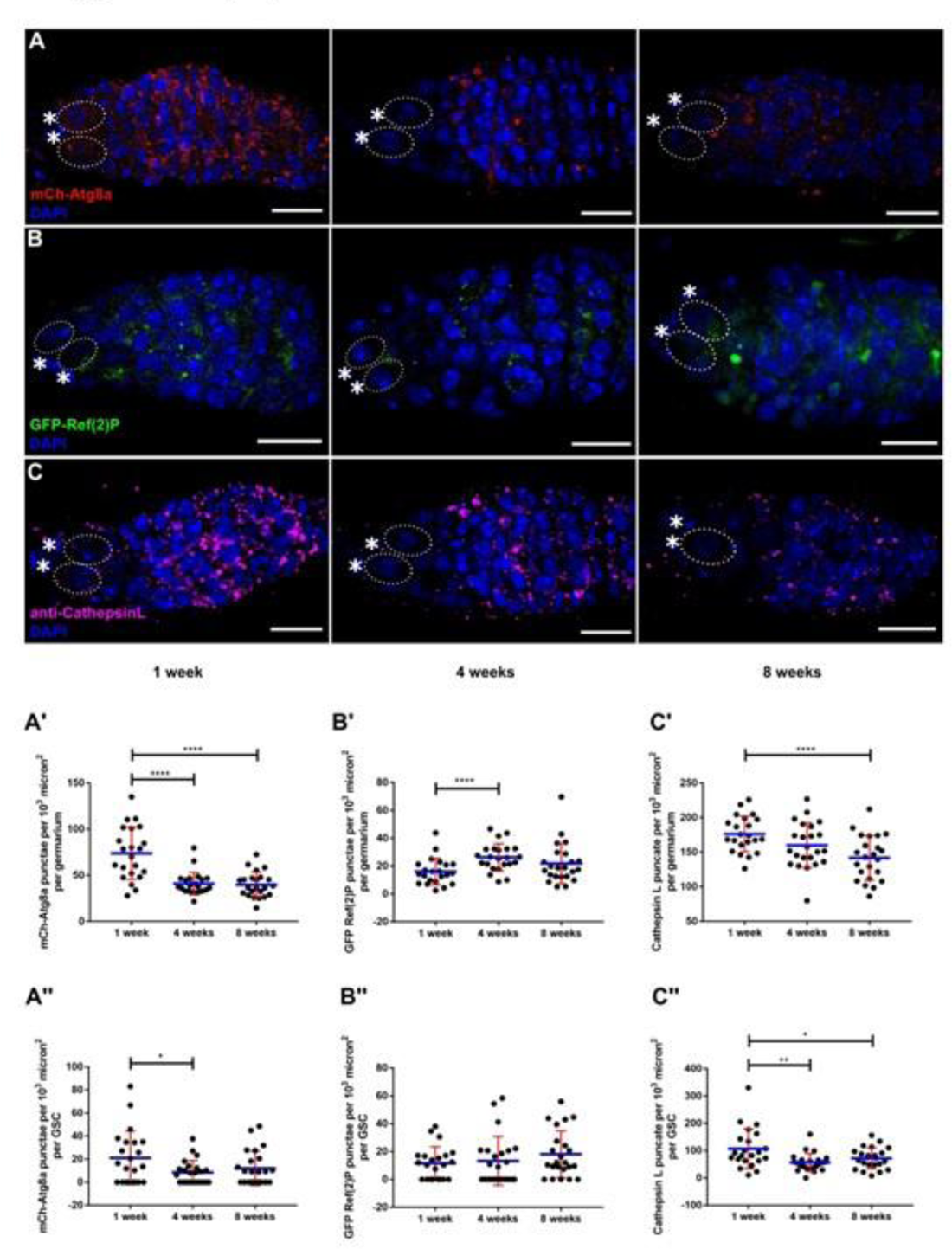
Autophagic flux reduces with age (A-C) Representative images for wildtype germaria (ywhsFLP1; nosP mCherry Atg8a; nosP GFP Ref(2)P) depicting mCherry-Atg8a positive puncta in red (autophagosomes) (A), GFP-Ref(2)P positive (cargo) in green (B) and CathepsinL (C) positive puncta in magenta (autolysosomes) at 1 week, 4 weeks and 8 weeks. Dotted ovals mark the GSCs and asterisk mark the cap cells. Nuclei are marked in blue. Scale bar-10µm. (A’-C’’) Interleaved scatter graph representing decrease in autophagosomes (A’-A’’), GFP Ref(2)P accumulation (B’-B’’) and decrease in autolysosomes (C’-C’’) in germarium and GSCs at 1 week, 4 weeks and 8 weeks respectively. Error bars represent standard deviation (S.D.) in red and blue lines represent mean. n=20. Genotype: ywhsFLP1; nos mCherryAtg8a/+; nos GFP Ref(2)P/+ *p < 0.05, **p < 0.01, ****p < 0.0001

**Supplementary figure 2.**
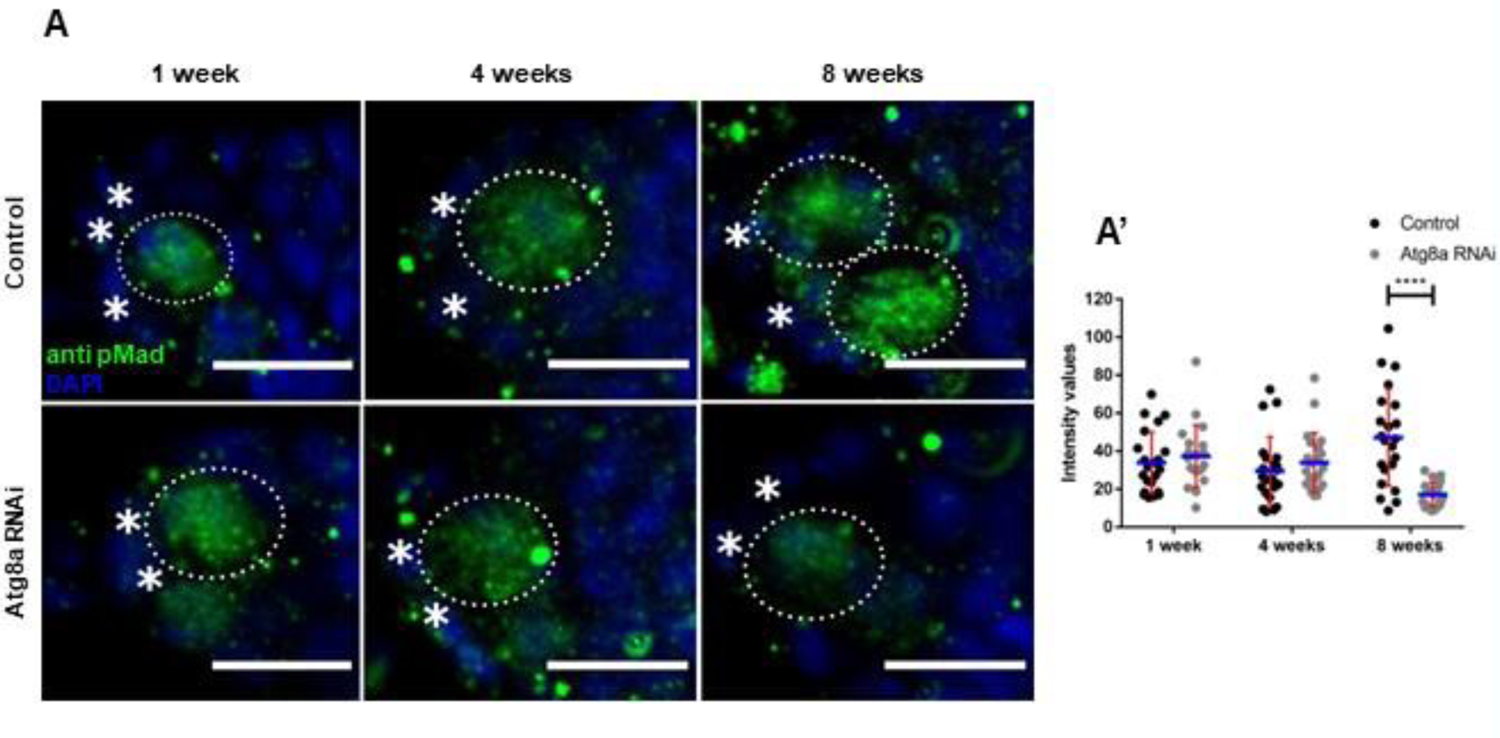
Effect of Atg8a knockdown on pMad levels (A) pMad in the GSCs (green) in controls (Atg8a RNAi/+) (top) and test (Atg8a RNAi/nos-Gal4) (bottom) at 1-8 weeks. Nuclei are marked in blue. Dotted ovals mark the GSCs and asterisk mark the cap cells. Scale bar-5µm. (A’) pMad intensities declines only at 8 weeks upon knockdown of Atg8a. Error bars represent standard deviation (S.D.) in red and blue lines represent mean, n=20. Control: ywhsFLP1/w; Atg8a RNAi/+; Atg8a RNAi/+, Atg8a RNAi: ywhsFLP1/ywhsFLP12; Atg8a RNAi/+; Atg8a RNAi/nos-Gal4. ****p < 0.0001

**Supplementary figure 3.**
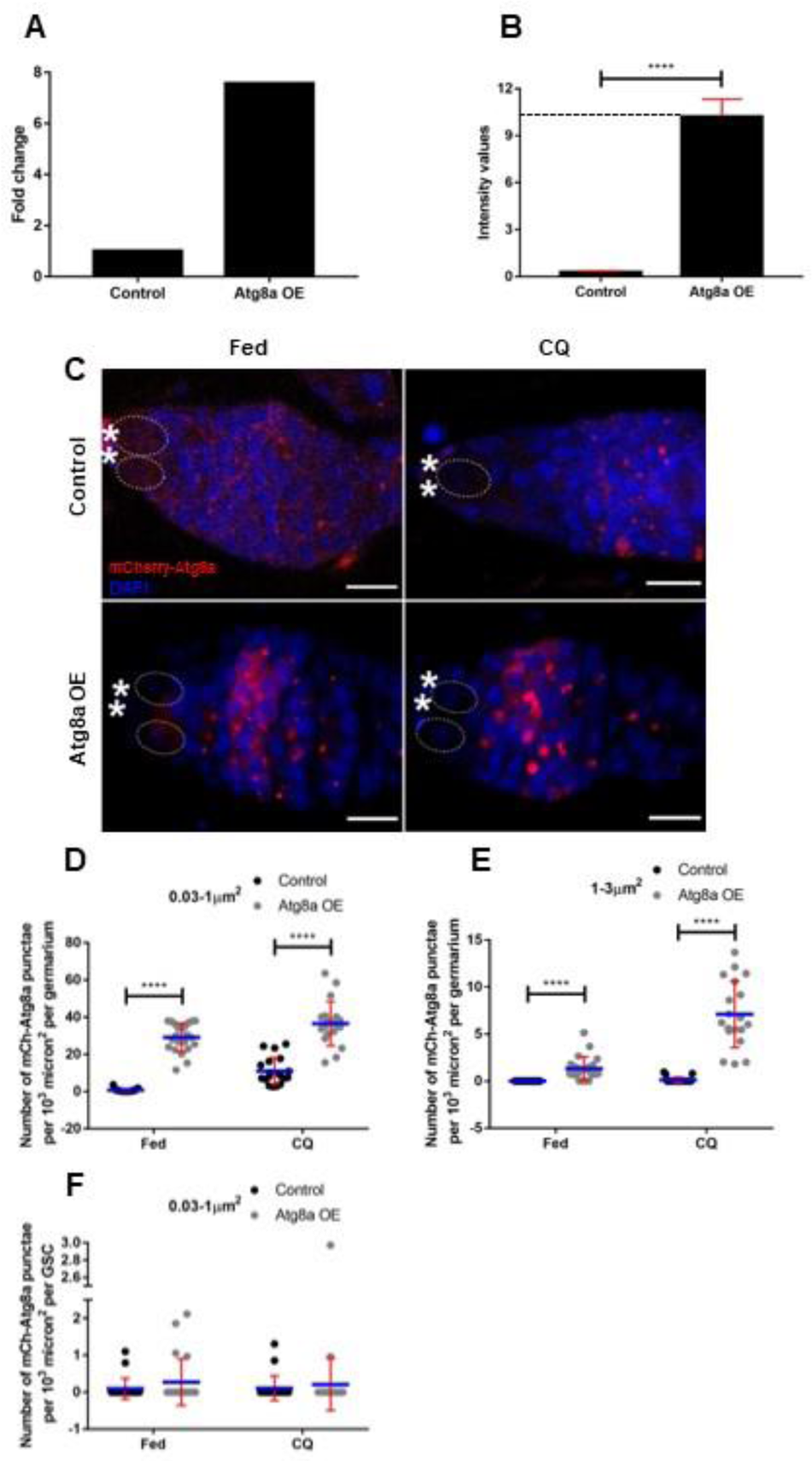
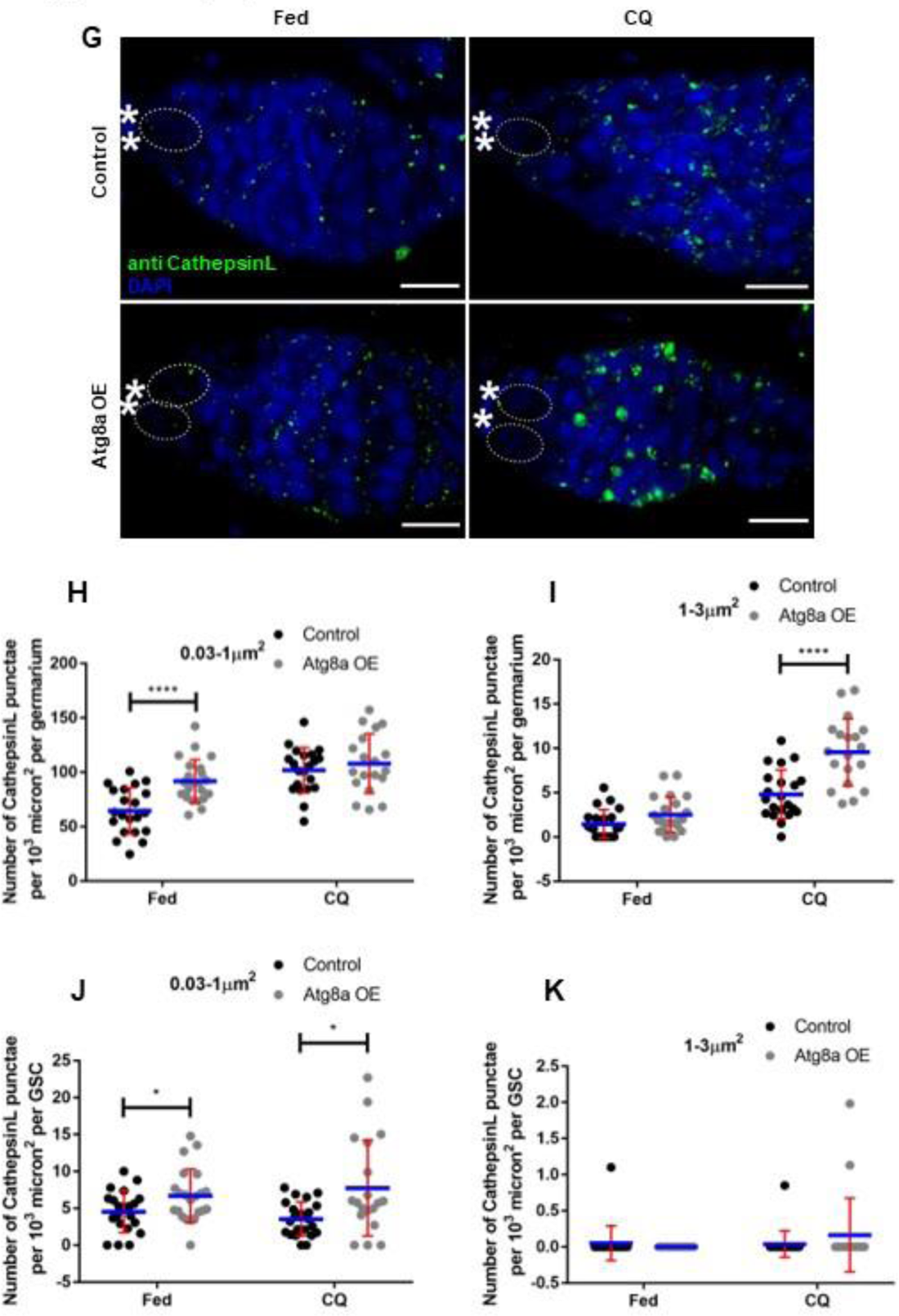
Atg8a affects size and number of autophagic vesicles (A) qPCR analysis shows 75 percent increase in Atg8a expression in test germaria as compared to controls. (B) Immunofluorescence intensity of mCherry in Atg8a OE shows 10 fold increase as compared to controls, n=12. (C) Representative images for indicated genotype fed on yeast and chloroquine (left to right) depicting autophagosomes (red). Nuclei are marked in blue. Dotted ovals mark the GSCs and asterisk mark the cap cells. Scale bar-10µm. (D-F) Interleaved scatter graph representing number of mCherry Atg8a puncta in germarium and GSCs. A size cut off (0.03-1µm^2^) and (1-3µm^2^) represents smaller and larger sized autophagosomes respectively. Increase in number of smaller as well as large sized autophagosomes in both fed and CQ treatment in germarium with Atg8a overexpression (D-E). Marginal increase in smaller sized autophagosomes observed in GSCs of test flies (F). Error bars represent standard deviation (S.D.) in red and blue lines represent mean, n=20. (G)Representative images for indicated genotype fed on yeast and chloroquine (left to right) depicting autolysosomes (green). Nuclei are marked in blue. Dotted ovals mark the GSCs and asterisk mark the cap cells. Scale bar-10µm. (H-K) Interleaved scatter graph representing number of CathepsinL puncta in germarium and GSCs. A size cut off of (0.03-1µm^2^) and (1-3µm^2^) represents smaller and larger sized autolysosomes respectively. Increase in number of smaller sized autophagosomes in both fed and CQ treatment in germarium (H) and GSCs (J) with Atg8a overexpression. Enlarged autolysosomes observed only upon CQ treatment in germarium with Atg8a overexpression (I). Number of larger sized autolysosomes do not change upon Atg8a overexpression in GSCs (K). Error bars represent standard deviation (S.D.) in red and blue lines represent mean, n=20. (A-B) Control: yw; UASp mCherry Atg8a/+, Atg8a OE: yw/w; UASp mCherry Atg8a/+; nos-Gal4/+. (C-K) Control: yw;UASp mCherry Atg8a/nosmCherry Atg8a;+, Atg8a OE: yw/w;UASp mCherry Atg8a/nosmCherry Atg8a; nos-Gal4/+. *p < 0.05, ****p < 0.0001

**Supplementary figure 4.**
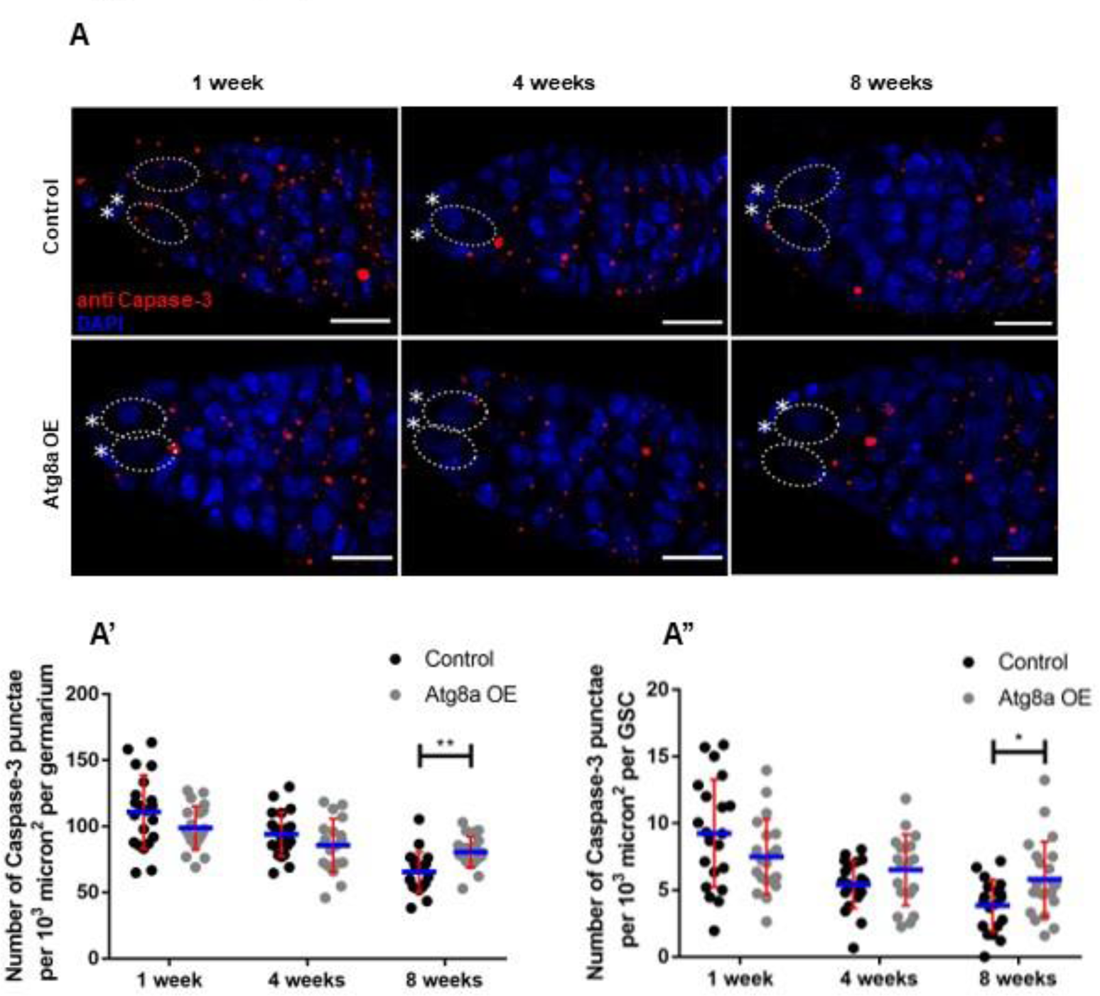
Increased autophagy lead to autophagy induced cell-death at 8 weeks. Distribution of caspase-3 puncta (red) in controls (UASp mChAtg8a/+) (top) and test (UASp mChAtg8a/nos-Gal4) (bottom) at 1-8 weeks. Nuclei are marked in blue. Dotted ovals mark the GSCs and asterisk mark the cap cells. Scale bar-10µm. (A’-A’’) Increase of cell death in 8 week old germaria (A’) and GSCs (A’’). Error bars represent standard deviation (S.D.) in red and blue lines represent mean, n=20. Control: yw;UASp mCherry Atg8a/+, Atg8a OE: yw/w;UASp mCherry Atg8a/+; nos-Gal4/+ *p < 0.05, **p < 0.01

**Supplementary figure 5.**
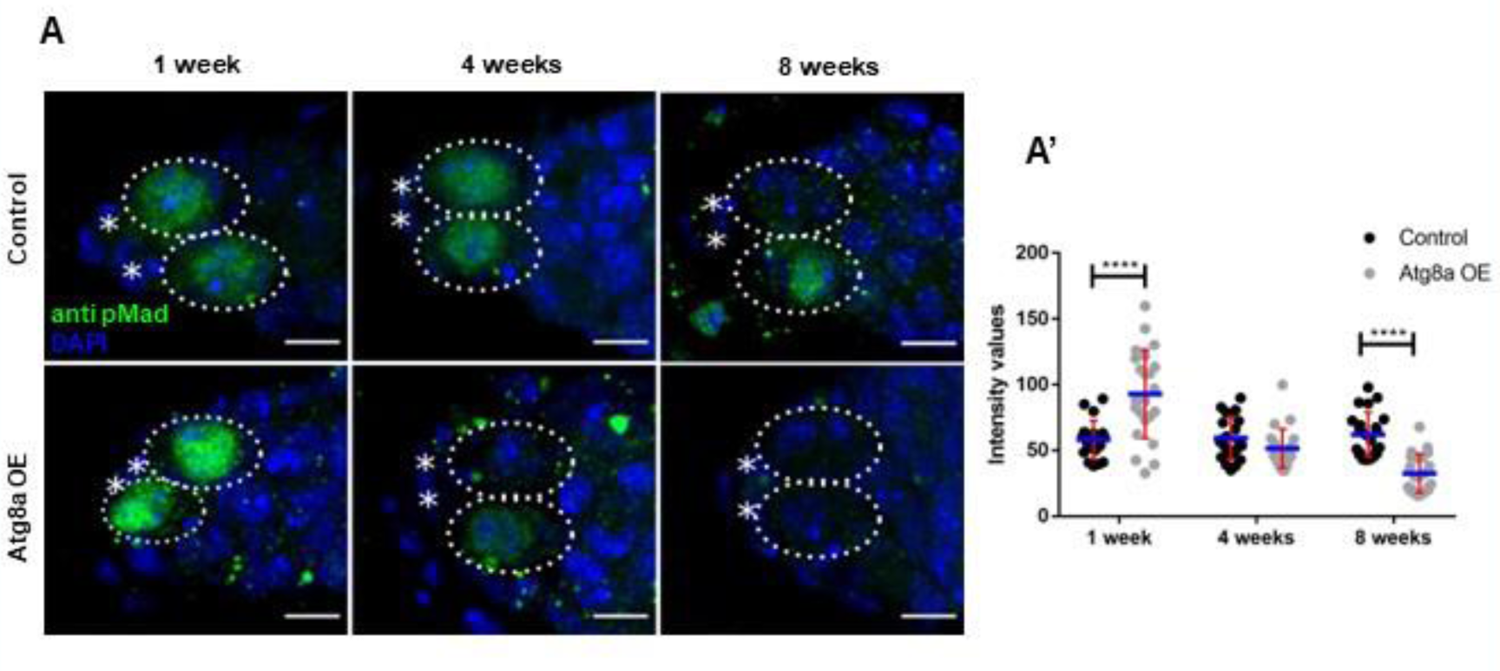
Effect of Atg8a overexpression on pMad levels pMad in the GSCs (green) in controls (UASp mChAtg8a/+) (top) and test (UASp mChAtg8a/nos-Gal4) (bottom at 1-8 weeks). Nuclei are marked in blue. Dotted ovals mark the GSCs and asterisk mark the cap cells. Scale bar-5µm. pMad intensities increase at 1 week and decrease at 8 weeks upon overexpression of Atg8a (B). Error bars represent standard deviation (S.D.) in red and blue lines represent mean, n=20, Control: yw;UASp mCherry Atg8a/+, Atg8a OE: yw/w;UASp mCherry Atg8a/+; nos-Gal4/+ ****p < 0.0001

## Author Contributions

NM and BVS designed the experiments. NM performed the experiments. Both authors wrote the manuscript and commented on it.

## Acknowledgments

We would like to thank Ms. Amruta Nikam for assisting with fly food preparation. Kiran Nilangekar for help with setting up qPCR. We acknowledge the confocal facility at ARI for assistance with imaging. Thanks to members of the Shravage lab for helpful discussions. We would like to thank Developmental Studies Hydridoma Bank, United States, for providing antibodies and plasmid constructs. Thanks to Prof. L. S. Shashidhara and IISER Fly community, for providing fly stocks. We would also like to thank Dr. P. K. Dhakephalkar, Director, Agharkar Research Institute, Pune, and entire ARI fraternity for support and access to facilities.

